# Ischemic stroke induces staged translational remodeling in mouse cortex independent from transcriptional responses

**DOI:** 10.1101/2025.08.15.670488

**Authors:** Sherif Rashad, Yuki Kitamura, Tomohito Nagai, Daisuke Ando, Abdulrahman Mousa, Hajime Ikenouchi, Hidenori Endo, Kuniyasu Niizuma

**Author notes:** **Equal contribution** (Sherif Rashad, Yuki Kitamura). **Corresponding author**: **Sherif Rashad, MD, PhD,**.

## Abstract

Ischemic stroke imposes acute oxidative and metabolic stress, yet how this reshapes mRNA translation remains poorly defined. We generated a time-resolved RNA-seq and ribosome-profiling atlas of mouse ischemic cortex at 1, 6 and 24 hours. Stroke produced early discordance between mRNA abundance and ribosome occupancy, revealing translational efficiency changes not captured by transcriptomics. The hyperacute phase was marked by a transient shift toward G/C-ending codon usage and a burst of 3′ untranslated region ribosome footprints consistent with stop-codon readthrough-like termination stress. Machine-learning analyses linked this response to stop-proximal ribosome queuing, 3′ untranslated region frame usage, local RNA structure and transcript architecture rather. From 6 to 24 hours, ischemic cortex developed progressive A-site pausing and altered coding-frame occupancy. Hidden Markov modeling localized a subset of frame-disruption events to discrete coding-sequence windows and machine learning revealed reproducible codon, amino-acid, nucleotide-composition and RNA-structure features associated with frameshifting. ORF-level analyses showed that many frame changes coincided with differential open reading frame usage. Late injury was characterized by increased upstream open reading frame engagement and 5′ untranslated region ribosome occupancy, inversely correlating with coding-sequence translation. These data identify staged translational remodeling after stroke that could not be captured via traditional mRNA or protein expression analysis.

## Introduction

Ischemic stroke (IS) remains one of the leading causes of long-term disability and mortality worldwide(1). Although rapid advances in reperfusion therapies over the past two decades have markedly improved patient outcomes(2,3), the global incidence of IS continues to rise, particularly in low– and middle-income countries(4). The pathophysiology of ischemic stroke is highly dynamic and involves a coordinated yet complex interplay among neurons, microglia, astrocytes, endothelial cells, and other brain-resident cell types(5). Neuronal death after ischemia is driven predominantly by excitotoxicity, oxidative stress, and microglia-mediated neuroinflammation(5,6), While regenerative approaches are being actively pursued(7), there remains an urgent unmet need for neuroprotective strategies capable of preserving neuronal integrity and preventing irreversible tissue loss(5,8). The limited success of prior neuroprotective interventions is often attributed to an incomplete understanding of the molecular programs engaged during ischemia.

Despite extensive transcriptomic work, including bulk and single-cell RNA-seq studies(9–11), our understanding of the translatome in stroke remains severely underdeveloped. This represents a major gap, as accumulating evidence shows a pronounced uncoupling between mRNA abundance and protein synthesis during acute cellular stress(12). Technologies that directly quantify ribosome-engaged mRNAs, such as ribosome profiling (Ribo-seq), offer a far more accurate representation of the cellular proteome and functional state compared to gene-expression–centric approaches(13,14). Yet, remarkably, Ribo-seq has never been applied to the study of ischemic stroke, leaving the translational landscape of injured neurons essentially unexplored.

In parallel, oxidative stress activates extensive epitranscriptional and post-transcriptional regulatory programs that modulate both mRNA translation and tRNA function(12,15). The tRNA epitranscriptome has emerged as a potent regulator of codon decoding and selective translation during stress(12,15–19). We previously demonstrated that ischemia induces widespread tRNA cleavage and production of tRNA-derived fragments (tDRs)(20,21). hinting at a deeper layer of translational and epitranscriptomic remodeling during stroke. However, no studies to date have systematically integrated translatome profiling with these regulatory processes in vivo.

Here, we address this critical knowledge gap by performing the first temporal Ribo-seq analysis in a mouse model of ischemic stroke. Our findings uncover multiple layers of translational deregulation, spanning codon-biased translation, stress-induced ribosomal reprogramming, stop-codon readthrough (SCRT), and frameshifting, that collectively shape the cortical brain response to ischemia. By defining these translational adaptive and maladaptive mechanisms, our work provides previously inaccessible insights into stroke biology and highlights new molecular pathways that may be leveraged for the development of next-generation neuroprotective therapeutics.

## Methods

### ➢ Animals

Distal middle cerebral artery occlusion (dMCAO) was induced on the left side in 12-week-old, 20-25 g male C57BL/6JJmsSlc mice as previously described(22). Animals were bought from a local vendor (Kumagai-Shoten Sendai) and housed in a controlled environment with a 12 h light/dark cycle and temperature regulated at 23°C. Animals were allowed to acclimatize for at least 1 week before surgery. All experiments were conducted according to protocols approved by the animal care facility of Tohoku University (Ethical approval number 2024-Ikoudou-005-01), the Animal Research: Reporting of In Vivo Experiments guidelines, and the Code of Ethics of the World Medical Association.

### ➢ dMCAO surgery

The permanent dMCAO model was performed according to previous reports(22,23). Briefly, mice were anesthetized with 2% isoflurane (30% oxygen, 70% nitrous oxide). Rectal temperature was maintained at 3°C using a heating pad with feedback control during surgery. The temporalis muscle between the left eye and ear was retracted along a 4-mm skin incision. A small bone window (∼1 mm diameter) was drilled to expose the distal MCA. The MCA was cauterized using bipolar electrocoagulation distal to the lenticulostriate branches. The incision was closed with 6-0 nylon sutures. Animals were monitored in a cage placed on a 3°C –heating plate for 1 h during anesthesia recovery. The left common carotid artery (CCA) was temporarily occluded with 6-0 nylon immediately before MCA cauterization and was released after 1 h.

Animals were euthanized at 1 h, 6 h, or 24 h after MCA cauterization. Sham-control animals underwent the same surgical procedure without MCA cauterization and CCA ligation and were euthanized after 1 h.

### ➢ TTC stain

At the designated time points after induction of dMCAO, mice were euthanized using isoflurane overdose inhalation followed by trans-cardiac perfusion with ice-cold saline to flush out the blood. Brains were dissected and 2-mm coronal sections of the brains were cut on a brain matrix and stained with 0.1% 2,3,5-triphenyltetrazoliumchloride (TTC; Sigma, Cat# T8877) dissolved in normal saline at 37°C for 15 min. Images were acquired using a flat-head scanner.

### ➢ Western blotting

At the designated time points, animals were sacrificed and trans-cardiac saline perfusion performed. Brains were dissected and the infarct area extracted and homogenized in T-PER tissue protein extraction reagent (Thermo Fisher, Cat# 78510) containing cOmplete protease inhibitor cocktail (Roche, Cat# 4693116001) and PhosSTOP phosphatase inhibitor (Roche, Cat# 4906837001). After centrifugation, the proteins were isolated from the supernatant and measured using Pierce BCA Protein Assay Kits (Thermo Fisher Scientific, Cat# 23227). Next, on Mini-PROTEAN TGX Precast Protein Gels (Bio-Rad, Cat# 4561096), equal protein loads were separated and then transferred to Trans-Blot Turbo Mini 0.2 µm PVDF Transfer Packs (Bio-Rad, catalog #1704156). Afterward, membranes were blocked with 5% skim milk (Fujifilm Wako, Cat# 190-12865) in phosphate-buffered saline with Tween 20 (PBS-T) (Takara, Cat# T9183). The primary antibody was then incubated overnight at 4°C, followed by a one-hour incubation at room temperature with Western BLoT Rapid Detect v2.0 (Takara, Cat# T7122A). Signal detection was performed using Pierce ECL substrate reagent (Thermo Fisher, Cat# 32106) on a ChemiDoc Touch system (Bio-Rad). The detected bands were analysed using Image Lab Software (BioRad). Beta-actin was used as a loading control. To reprobe the membrane, Restore stripping buffer (Thermo Fisher, Cat# 21059) was utilized.

#### List of antibodies used

beta-Actin Rabbit Monoclonal (1:3000, Cell signaling, Cat# 4970).

eRF1 Rabbit Polyclonal (1:1000, Invitrogen, Cat# PA5-28777)

Sephs2 Rabbit Polyclonal (1:1000, Invitrogen, Cat# PA5-27950)

Doublecortin (DCX) Rabbit Polyclonal (1:500, Abcam, Cat# ab18723)

Smad2 Rabbit Polyclonal (1:1000, Invitrogen, Cat# PA5-88144)

### ➢ Immunofluorescence staining

Mice were euthanized by aspirating excess isoflurane at the designated timepoints after dMCAO procedure. Mice were transcardially perfused with saline to remove the blood and subsequently with 4% paraformaldehyde in phosphate buffer (Nakarai Tesque, Cat# 09154-85). The fixed middle third of the brain, including the ischemic region, was removed and embedded in OCT compound (Sakura Finetek, Cat# 4583) to be frozen by liquid nitrogen. The prepared brain was then sectioned at a 10 μm thickness with a cryostat. The sections were washed with 1 x PBS, incubated with blocking buffer with 0.8% Block Ace (UKB, Cat# UK880), 1% BSA (Fujifilm Wako, Cat# 010-25783), and 0.3% Triton X-100 (Nacalai Tesque, Cat# 35501-15) in PBS for 1 h at room temperature before adding primary antibodies (details below) diluted in blocking buffer, and incubated overnight at 4 °C. Subsequently, they incubated with conjugated secondary antibodies of AlexaFluor 568 (1:500; Invitrogen, Cat# A11041, A10042, and A10037) and AlexaFluor 647 (1:500; Invitrogen, Cat# A21244, A21136, and 21244) in antibody-dilution buffer with 0.2% Block Ace, 0.2% BSA and 0.3% Triton X-100 in PBS and DAPI (1:500; Invitrogen, Cat# D1306) was used for counterstaining. Samples were inspected under a laser confocal microscope (OLYMPUS, FV3000).

#### Primary antibodies used

Iba-1 Rabbit Polyclonal (1:500, Fujifilm Wako, Cat# 019-19741)

NeuN Rabbit Monoclonal (1:1500, Abcam, Cat# ab177487)

GFAP Rabbit Monoclonal (1:200, Invitrogen, Cat# 130300)

### ➢ Ribo-seq library preparation

Ribo-seq was conducted as previously described(24,25). Briefly, at the designated time point, animals were sacrificed and perfused using ice-cold PBS containing 100 µg/ml cycloheximide (Sigma, Cat# 01810), followed by extraction of the brain. Brains were extracted and sectioned into three equal parts in the sagittal plane. The leftmost third of the brain, corresponding to Bregma –2 mm to +2 mm, where the infarct occurs, was immediately snap-frozen in liquid nitrogen. Dissection was conducted on ice, and all tools were decontaminated with RNase AWAY(Thermo Fisher, Cat# 7002PK) prior to use. Tissues were homogenized in polysome lysis buffer (20 mM Tris-Cl, pH 7.5, 150 mM NaCl, 5 mM MgCl_2_, 1 mM dithiothreitol (Sigma, Cat# D9779), 100 µg/ml cycloheximide, and 1% Triton X-100) and centrifuged at 13000 x g at 4°C for 15 min. Ribosome foot-printing was performed on the supernatant collected by adding 1.25 U/μl RNase I (NEB, Cat# M0243) to 500 μl clarified lysate and incubating samples on a rotator mixer for 45 min at room temperature. QIAzol (QIAGEN, Cat# 79306) reagent was added, and RNA was extracted using miRNeasy Mini Kit (QIAGEN, Cat# 217004). Ribosome-protected fragments (RPFs) were selected by isolating RNA fragments of 27-35 nucleotides (nt) using TBE-Urea PAGE gel. rRNA depletion was conducted using NEBNext rRNA depletion kit v2 (NEB, Cat# E7400) followed by end-repair using T4 PNK after purification of the rRNA depleted samples using Oligo Clean & Concentrator Kits (Zymo Research, Cat# D4060). The preparation of sequencing libraries for ribosome profiling was conducted via the NEBNext Multiplex Small RNA Library Prep kit for Illumina according to the manufacturer’s protocol (NEB, Cat# E7300S). Library quality was assessed on an Agilent Bioanalyzer 2100 using the Agilent DNA 1000 kit (Cat# 5067-1504). Pair-end sequencing reads of size 150 bp were produced for Ribo-seq on the Illumina HiSeq X-ten system. Four biological replicates were prepared per group.

### ➢ Ribo-seq data analysis

Quality control for Raw Fastq was performed using FastQC. Next, adapter trimming and collapsing the pair-end reads into one was done using Seqprep. Trimmomatic(26) was used to further clean low-quality reads. Bowtie2(27) was used to align the reads to a reference of rRNA and tRNA genes (mm10) to remove contaminants. After that, reads were aligned to the genome (mm39 downloaded from UCSC), and reads counted using FeatureCounts(28) with the tag “-t CDS” to generate counts of protein coding genes only. RiboWaltz(29) was used for metagene analysis, P-site offset determination, and quality control of Ribo-seq samples. Pausepred offline was used for ribosome pause sites determination (https://github.com/romikasaini/Pausepred_offline). Visualization of Pausepred results was performed in *R* (version 4.5.0).

### ➢ RNA-seq library preparation

The ischemic brain regions described in Ribo-seq section, or anatomically matched areas from sham animals, were lysed using QIAzol Lysis Reagent (QIAGEN, Cat# 79306), and total RNA was extracted with the miRNeasy Mini Kit (QIAGEN, Cat# 217004) according to the manufacturer’s instructions. RNA purity and concentration were measured using a NanoDrop spectrophotometer (Thermo Fisher, Cat# ND-ONE-W), and RNA integrity was assessed using the RNA 6000 Nano Kit (Agilent, Cat# 5067-1511) on the Agilent 2100 Bioanalyzer. Only samples with an RNA Integrity Number (RIN) of 7 or higher are used for RNA sequencing (RNA-seq).

RNA-seq was performed using three biological replicates per group, with mRNA enriched using the NEBNext Poly(A) mRNA Magnetic Isolation Module (NEB, Cat# E7490) and libraries prepared using the Ultra II Directional RNA Library Prep Kit for illumina (NEB, Cat# E7760) according to the manufacturer’s instructions. Library quality was assessed on an Agilent Bioanalyzer 2100 using the Agilent DNA 1000 kit (Cat# 5067-1504), and library concentrations were quantified with the NEBNext Library Quant Kit for Illumina (NEB, Cat# E7630). Sequencing was conducted on an Illumina NovaSeq X Plus platform using 150 bp paired-end reads. Raw Fastq files underwent quality control with FastQC, followed by trimming with Trimmomatic(26) to remove adapter sequences and low-quality reads. Clean reads were aligned to the Mus musculus (mm39) reference genome (UCSC) using the splice-aware aligner HISAT2(30), and gene-level counts were generated with FeatureCounts(28) using UCSC Refseq annotation.

### ➢ Differential gene expression and translational efficiency analysis

Differential gene expression and translational efficiency analysis were performed in *R* using DESeq2(31) and (https://github.com/smithlabcode/riborex) with Limma-voom based approach respectively. Pathway enrichment analysis, correlation analysis, and codon analysis of sequencing datasets were all performed using *R* language programming.

### ➢ GC3 and isoacceptors codon analysis

Isoacceptors frequency and total codon frequency were analyzed as we previously reported(24,25,32,33). We used the protein coding transcript fasta from Gencode (mouse V37) to extract the coding sequences. We selected the longest transcript per gene as representative transcript for codon analysis. We used the R package coRdon to generate codon tables.

GC3 scores for each gene were calculated using this formula:

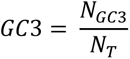

Where *N_GC3_* is the number of codons ending with G or C in the mRNA sequence and *N_T_* is the total number of codons in the mRNA sequence. The GC3 score varies from 0 (no G/C-ending codons) to 1 (All codons are G/C-ending).

Isoacceptors codon frequencies were calculated as follows:

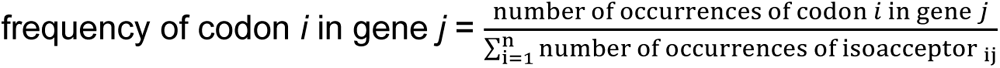

where isoacceptors refers to the synonymous codons for the amino acid encoded by codon *i* and n refers to the number of isoacceptors for a given amino acid (2 ≤ n ≤ 6). The isoacceptors scores ranged from 0 to 1, representing the ratio of each synonymous codon in the mRNA sequence.

The t-statistic (T-stat) describing the codon frequency was calculated as:

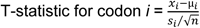

Where x*_i_* refers to the mean frequency for codon *i* in the sample, µ*_i_* refers to the mean frequency for codon *i* across the Mus musculus genome, s*_i_* refers to the standard deviation of the frequency for codon *i* in the sample, and n refers to the sample size.

T-statistics value of > 2 or < –2 indicates statistical significance (*p* < 0.05) versus the background (i.e. all genes)

### ➢ Gene level frameshifting analysis

To analyze frameshifted RPFs, we created a custom python script that first estimates the P-site offset using the same logic of RiboWaltz(29). After P-site offset calculation per RPF length for each bam file, reads were counted and mapped to the P-site to determine their frame, based on their unique P-site offset in each bam file, whether it is 0, +1, or +2 frame. Frameshift Score (FS) for a gene was calculated as:

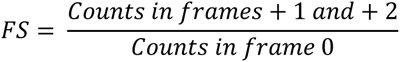

Thus, FS > 1 means frameshifting for a specific gene, while FS < 1 means in-frame reads for a gene.

After calculating FS scores and RPF counts for all samples, genes with total CDS RPF ≥ 20 in at least 3 biological replicates per group were retained for analysis and the results merged into a single matrix. Limma-Trend(34) was used to compare between each condition and the sham group, with significance defined as |log2FC| ≥ 1 and FDR ≤ 0.05.

#### Identification of ribosomal frameshift loci from Ribo-seq data

To identify sites of aberrant translational frameshifting, we developed a computational framework that quantifies local deviations from the expected 0-frame periodicity in gene with statistically significant frameshifting in stroke conditions versus sham. For each candidate transcript, P-site counts were aggregated by codon and frame across biological replicates within each group. Codon-level frame profiles were modeled using a three-state HMM, with states corresponding to canonical frame 0, +1 frame, and −1 frame relative to the annotated CDS. HMM emissions were multinomial over the three frame-count classes, and the Viterbi path was used to assign the dominant frame state at each codon. Candidate loci were defined as sustained off-frame Viterbi runs of at least six codons.

For each candidate seed, we extracted a local codon-space window comprising 10 codons upstream and 10 codons downstream of the six-codon minimal off-frame seed, corresponding to a 26-codon/78-nt window unless clipped by CDS boundaries. Candidate windows were required to contain ≥30 RPFs and to show significant differential frame usage between stroke and sham by a 3×2 G-test at α = 0.05. Genes entering the locus-calling step were required to have ≥20 CDS RPFs per group. For transcripts with multiple candidate loci, ±90 nt around the first locus was masked before searching for additional loci. Direction labels were assigned in transcript/coding-frame coordinates, with HMM state 1 corresponding to +1 and state 2 corresponding to −1, independent of genomic strand.

Because no ground-truth catalogue of stroke-induced mammalian frameshift loci is available, the HMM was used as an approximate localization procedure rather than as definitive proof of ribosomal slippage at a single nucleotide. Thresholds were selected to prioritize sustained and coverage-supported frame changes: a minimum six-codon off-frame run, ≥30 RPFs in the candidate window, and significant differential frame usage by G-test. We therefore interpret HMM-positive loci as a high-confidence subset of gene-level frameshift events. Genes with significant gene-level FS changes but no discrete HMM locus were retained as diffuse frame-disruption candidates and were considered separately from localized loci.

#### Sequence-context characterization of frameshift loci

For each identified locus, we extracted a fixed-length, strand-aware RNA window centered on the frameshift event. We quantified known frameshift-promoting features, including slippery heptamers (XXXYYYZ), homopolymeric A/U tracts, codon runs (e.g., poly-Pro, poly-Lys), and viral-like motifs. Local secondary structure was analyzed using minimum free energy (MFE), ensemble free energy, and MFE frequency computed by RNAfold(35). GC/AU composition, CpG frequency, nucleotide entropy, and G-quadruplex-forming sequences were extracted. Positional features (relative location within CDS, exon boundaries, and codon span) and amino-acid–level patterns (poly-Pro, poly-basic tracts, acidic–proline transitions) were quantified.

We quantified local nucleotide diversity using Shannon entropy of base composition in the FS-centered window (H = −∑ pᵢ log₂ pᵢ for i ∈ (A,U,G,C)). Lower entropy reflects strong compositional bias (e.g. AU-rich or GC-rich tracts), whereas higher entropy indicates compositionally balanced sequences. Frameshift loci tended to reside in windows with modestly reduced entropy compared to matched background windows, consistent with enrichment of homopolymeric tracts and biased sequence contexts.

#### Mapping frameshift events to transcript structures and functional consequences

To determine the consequences of each frameshift event on coding output, genomic coordinates were mapped to transcript-specific CDS coordinates using a GTF annotation. For each gene, all annotated transcripts containing a CDS were considered. A transcript was selected preferentially if (i) it matched the transcript ID carried from the upstream analysis, or (ii) its CDS encompassed the frameshift coordinate. For the selected transcript, we reconstructed its CDS and 3′UTR sequence directly from the genome. Frameshifted open reading frames (ORFs) were simulated by applying a +1 or −1 shift at the exact nucleotide where the out-of-frame state was detected. We determined:

- **PTC (premature termination codon)** if the shifted ORF terminated upstream of the annotated stop codon;
- **SCRT (stop-codon readthrough–like extension)** if the shifted ORF terminated downstream of the annotated stop;
- **NMD susceptibility** if the PTC occurred >50 nt upstream of the last exon–exon junction in the CDS.

The canonical and frameshifted CDS and protein sequences were generated for all events and classified into these functional categories.

#### Machine-learning analysis of frameshift loci

For each HMM-localized frameshift locus, we extracted a strand-oriented local coding-sequence window centered on the inferred frameshift signal and quantified features corresponding to the same families used for downstream TreeSHAP interpretation. The primary window consisted of the six-codon HMM off-frame seed plus 10 codons upstream and 10 codons downstream, corresponding to 26 codons/78 nt unless clipped by CDS boundaries. Features were computed in transcript/coding-frame orientation and included nucleotide composition, GC and GC3 fraction, CpG observed/expected ratio, Shannon nucleotide entropy, codon frequencies, amino-acid composition, hydrophobic amino-acid fraction, longest hydrophobic run, proline fraction, CDS-relative position, RNAfold-derived local structure metrics, predicted G-quadruplex sequence content, and predefined sequence/motif indicators.

To define negative controls, we generated CDS-matched background windows. The primary background consisted of non-overlapping windows sampled from the same transcript and matched to the local CDS context of each positive FS window, while excluding regions overlapping any HMM-localized FS locus. As a sensitivity analysis, we also generated global CDS background windows sampled from CDS regions outside the corresponding positive event set. Background construction and window-length distributions were recorded as QC outputs.

For all ML analyses, non-biological and potentially circular variables were excluded from predictor matrices, including gene/transcript identifiers, genomic coordinates, strand, timepoint labels, window-length/QC flags, HMM state variables, read-depth variables, and frame-usage statistics used during upstream locus calling. Stop-derived features, including local stop-codon fraction, stop-codon amino-acid features, and stop-codon triplet frequencies, were also excluded because they are better interpreted as predicted consequences of frame displacement rather than causal determinants of frameshifting. Missing numeric values were imputed using training-set medians, with non-finite columns removed where applicable.

We trained binary classifiers for five tasks at 6 h and 24 h: all FS loci versus matched background, FS-UP loci versus matched background, FS-DOWN loci versus matched background, FS-UP versus FS-DOWN loci, and +1 versus −1 FS loci. To avoid leakage among multiple windows from the same gene, all classifiers used gene-grouped 80/20 train/test splits in which all windows from a given gene were assigned exclusively to either training or held-out testing. Splits were repeated 100 times with independent random seeds. Model performance was evaluated on held-out test data using ROC-AUC, PR-AUC, and balanced accuracy.

Before final XGBoost interpretation, we compared elastic net logistic regression, random forest, and XGBoost classifiers across repeated gene-grouped splits. The final models used XGBoost with a binary logistic objective, max_depth = 4, eta = 0.05, subsample = 0.8, colsample_bytree = 0.8, and min_child_weight = 3. Class imbalance was handled using training-set-derived positive-class weights. The number of boosting rounds was selected within each training set by stratified cross-validation using up to 400 rounds and early stopping after 20 rounds. The final model for each iteration was then evaluated only on the held-out test set.

Feature importance was assessed using XGBoost gain and native TreeSHAP values computed on held-out test windows. For each task and timepoint, mean absolute TreeSHAP values were summarized across repeated splits and aggregated into biologically interpretable feature families. Codon-usage features included codon-frequency variables extracted from the FS-centered window. Amino-acid context features included amino-acid composition and derived peptide-context measures such as hydrophobic amino-acid fraction, longest hydrophobic run, and proline fraction. Nucleotide-composition features included GC/GC3/AU fractions, CpG observed/expected metrics, and nucleotide entropy. RNA-structure features included RNAfold-derived local structure metrics such as MFE, ensemble free energy, MFE frequency, and predicted G-quadruplex sequence content. Motif/tract features included predefined frameshift-associated sequence features such as homopolymeric A/U tracts, poly-amino-acid motifs, and amino-acid transition motifs. CDS-position features described the relative location of the window within the coding sequence. Sensitivity analyses included label-permutation null models, models excluding all CDS-position features, high-confidence HMM loci only, coverage-matched positive-only tasks, global CDS background models, and pooled H6/H24 models with timepoint as a covariate.

### ➢ Stop codon readthrough (SCRT) analysis

RiboWaltz-derived P-sites were used to quantify CDS and 3′UTR ribosome occupancy. RPFs of 25–36 nt were retained. For each transcript, the CDS stop coordinate and annotated 3′UTR length were obtained from the transcript annotation. In the primary analysis, 3′UTR P-sites were counted from a dynamic transcript-specific window beginning immediately downstream of the annotated stop codon and spanning the first 50% of the annotated 3′UTR, with a minimum window length of 30 nt and maximum length of 300 nt. The primary SCRT count model did not require 3′UTR frame locking. Genes were retained in a sample if they had ≥20 CDS P-sites, ≥1 3′UTR P-site, and ≥50 total CDS + 3′UTR P-sites. Genes were included in differential testing if they passed these thresholds in at least three samples within at least one group.

Per gene Ribosome Readthrough Score (RRS) was calculated as:

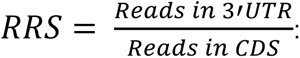

Differential SCRT was tested using a beta-binomial GLM comparing each stroke group with the sham group. The reported SCRT log2FC represents the log2 fold-change in the fitted 3′UTR ribosome proportion, UTR/(UTR+CDS), between stroke and sham. P values were adjusted using Benjamini–Hochberg FDR, and significant SCRT changes were defined as |log2FC| ≥ 1 and FDR ≤ 0.05. Metagene plots for selected genes were created using RiboWaltz, extending the plots to the 3’UTR as needed. GC3 and isoacceptors codon frequencies analysis was performed as explained above on significant genes.

#### Stop-Proximal Metagene Profiling and pre-stop ribosome occupancy calculation

To compare genes with significant changes in SCRT events in stroke vs control, we extracted the reads mapped to a window 60 nucleotides upstream of the stop codon. The window was binned in the codon space, to generate normalized P-sites at each codon site upstream of the stop codon. For the background genes, we selected a random set of genes with comparable CDS coverage to the foreground set to avoid inflation of pre-stop p-site density of the background genes due to large number of genes selected or differences in coverage.

Pre-stop ribosome occupancy was calculated 30 nucleotides (i.e. 10 codons) upstream of the stop codon. This metric was used for machine learning modeling.

#### 3′UTR Frame-Usage Analysis

To evaluate the dominant frame in the 3’UTR reads, we assigned a frame for each read based on the P-site offset calculated for the CDS. Frame-usage matrices were aggregated across all genes and timepoints to detect frame biases associated with stop codon readthrough (SCRT).

#### RNA Structural Features Around the Termination Codon

We extracted a window of 100 nucleotides downstream of the stop codon and calculated minimum free energy (MFE) using Vienna RNAfold(35).

#### Machine-learning-based identification of SCRT determinants

Machine-learning analyses were performed to identify transcript features associated with stroke-induced stop-codon readthrough (SCRT). Separate input tables were generated for each post-stroke timepoint (H1, H6, and H24). SCRT status was assigned from the differential SCRT beta-binomial model, with significant SCRT changes defined as |log2FC| ≥ 1 and FDR ≤ 0.05. Genes with increased SCRT were labeled SCRT-UP, genes with decreased SCRT were labeled SCRT-DOWN, and genes without significant SCRT changes were used as the non-SCRT background pool.

For each task, background genes were selected by coverage matching rather than by using all non-significant genes. Specifically, SCRT-positive genes were matched to non-SCRT background genes using nearest-neighbor matching on pre-stop mean CDS ribosome density, thereby reducing the possibility that classifiers learned trivial read-depth differences. For SCRT-UP versus SCRT-DOWN models, the smaller direction class was matched to genes from the opposite direction using the same coverage-matching strategy. Model-comparison analyses evaluated four classification tasks: all SCRT-altered genes versus matched background, SCRT-UP genes versus matched background, SCRT-DOWN genes versus matched background, and SCRT-UP versus SCRT-DOWN genes. Final XGBoost analyses focused on all SCRT-altered genes versus background, SCRT-UP genes versus background, and SCRT-UP versus SCRT-DOWN genes.

Five feature panels were used. The stop-neighborhood panel included transcript architecture features, global/stop-proximal sequence features, stop codon identity, pre-stop nucleotide-composition features, and identity of the five codons immediately upstream of the stop codon. The stop-neighborhood-no-stop panel was identical except that stop codon identity was removed. The full-context panel added ribosome-context features to the stop-neighborhood panel, and the full-context-no-stop panel removed stop codon identity from this full feature set. The ribosome-context-only panel included only frame-resolved proximal 3′UTR ribosome usage and pre-stop ribosome enrichment. Feature families were defined as follows: transcript architecture (log-transformed CDS length and 3′UTR length), global sequence (CDS GC3, stop-window GC content, and RNAfold-derived stop-window minimum free energy), stop identity (TAA/TAG/TGA identity), pre-stop composition (GC fraction, GC3 fraction, and nucleotide entropy across the five pre-stop codons), pre-stop codon identity (codon identities from −5 to −1 relative to the stop codon), and ribosome context (proximal 3′UTR frame 0/1/2 fractions and pre-stop ribosome enrichment).

Numeric and categorical predictors were processed separately. Numeric predictors were retained as continuous variables, whereas categorical predictors such as stop codon identity and pre-stop codon identities were one-hot encoded. Missing categorical values were encoded as a separate missing level. Missing or non-finite numeric values were imputed using medians calculated from the training set only and then applied to the held-out test set.

Elastic-net logistic regression, random forest, and XGBoost classifiers were first compared across timepoints, SCRT tasks, and feature panels using 50 repeated gene-grouped 80/20 train/test splits. Gene grouping ensured that each gene was assigned exclusively to either the training set or the held-out test set within each iteration. Elastic-net models were fit using cross-validated glmnet with alpha = 0.5, random forests were fit using 500 trees, and XGBoost models used a binary logistic objective with max_depth = 4, eta = 0.05, subsample = 0.8, colsample_bytree = 0.8, min_child_weight = 3, and training-set-derived scale_pos_weight.

For final interpretation, repeated XGBoost models were trained using the same gene-grouped 80/20 held-out strategy across 100 iterations. Final models used fixed 100 boosting rounds to maintain consistency across repeated runs and small SCRT task sizes. For each final model, label-permutation controls were generated using the same splitting and modeling workflow after permuting class labels. Model performance was summarized as held-out ROC-AUC, PR-AUC, and balanced accuracy across repeated iterations. Feature importance was assessed using XGBoost gain and native TreeSHAP values computed on held-out test genes. TreeSHAP values were summarized both for individual features and after aggregation into the feature families described above.

### ➢ ORF analysis

Translated open read frames (ORFs) were calculated using RiboCode(36). We fixed the start codon as ATG to reduce complexity and limited the ORFs to a minimum 6 codons for ORF identification. RibCode’s results were then imported into R, and additional filtering (minimum P-sites per ORF ≥ 10, minimum codons/amino acids ≥ 10, frame 0 P-site counts ≥ 50% of all ORF P-site counts, and RiboCode significance (*p* value) ≤ 0.05) were implemented prior to downstream analysis. Differential ORF expression was calculated using EdgeR(37) using ORF P-site counts as input. The canonical ORF for each gene was defined as the longest ORF with the highest 0 frame P-sites. ORFs designated as annotated and protein coding were primarily defined as canonical. If multiple annotated and protein coding ORFs were detected per gene, the longest with the highest P-site counts was selected as the canonical. The dominant translated ORF was determined by calculating the ratio of ORF P-site counts for all ORFs derived from the same gene and selecting the ORF that has a ratio of ≥ 60%. Differential ORF usage (DOU) was performed using DRIMSeq(38) using a Dirichlet–multinomial model.

#### Analysis of differential upstream translation

To analyze genes with changes in their 5’UTR to CDS translation ratio, we used the same approach employed in the SCRT analysis but rather than comparing CDS and 3’UTR P-site counts with GLM-betabinomial modeling, we compared CDS and 5’UTR P-site counts between stroke and sham groups. Significance level was set as |log2FC| ≥ 1 and FDR ≤ 0.05. Upregulation indicates more reads in the 5’UTR compared to CDS in the condition versus sham and vice versa.

## Results

### ➢ Temporal dynamics of translation reflect stroke pathophysiological evolution over time

To profile translational responses to stroke, we used the distal middle cerebral artery occlusion (dMCAO) model by occluding the cortical distal M4 branch of the MCA in male mice (Figure 1a). Cortical infarction was evident at 24 h (Figure 1b). We therefore sampled tissue at 1, 6, and 24 h to capture early, intermediate, and established injury states. Immunofluorescence of the infarct region showed a progressive loss of neurons and astrocytes in the core, whereas microglia were evident at 24 h (Figure 1c).

**Figure 1:**
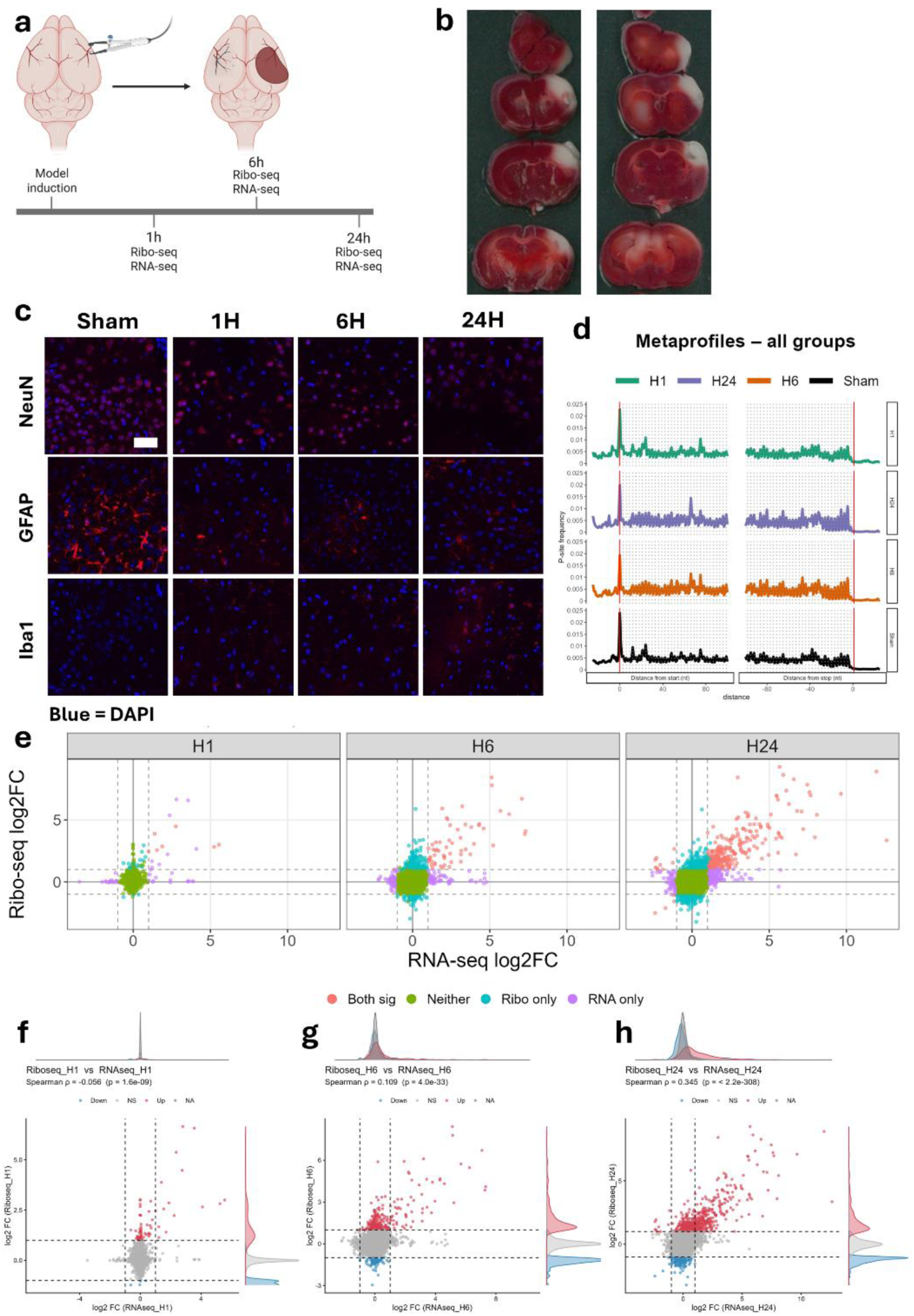
I**s**chemic **stroke leads to transcription-translation discordance. a:** Schematic showing the method for inducing stroke and the analysis performed at different timepoints after stroke. **b:** TTC stain showing the area of ischemic infarction 24 hours after dMCAO model induction. **c:** Immunofluorescence staining showing neurons (stained by NeuN), astrocytes (stained by GFAP), and microglia (stained by Iba1) after dMCAO model induction. bar = 50µm. **d:** Group-level metagene plot showing the cumulative 3-nucleotide periodicity indicating detection of translating ribosomes in the Ribo-seq analysis. **e:** Quadrant scatter plots showing the significant genes in the RNA-seq and Ribo-seq analyses at each time point. **f-h:** Spearman’s rank correlation analysis scatter plots showing the correlation between gene RNA-seq and Ribo-seq log2 fold change values at each time points (1 hour, 6 hours, and 24 hours respectively).

We next profiled molecular changes using matched RNA-seq and Ribo-seq at each timepoint. Ribo-seq libraries showed the expected 3-nt periodicity, confirming capture of translating ribosomes (Figure 1d). Stroke induced progressive changes in both mRNA abundance (RNA-seq) and ribosome occupancy (Ribo-seq) relative to sham controls (Figure 1e). Translational efficiency (TE; Ribo-seq normalized to RNA-seq, see Methods) was significantly altered at 1 h (320 genes; |log2FC| ≥ 1, FDR ≤ 0.05) and increased in intensity at later timepoints (927 genes at 6 h; 849 genes at 24 h) (Supplementary Figure 1a). Consistent with early transcription-translation decoupling, RNA-seq and Ribo-seq log2FC values were essentially uncorrelated at 1 h and 6 h (Figure 1f-g; Spearman’s ρ = −0.056 and 0.109, respectively), but became more aligned by 24 h, although still modestly (Figure 1h; Spearman’s ρ = 0.345). This suggests partial synchronization of transcriptional and translational programs as injury consolidates.

To interpret these temporal patterns, we performed pre-ranked GSEA using Gene Ontology Biological Process (GOBP) sets for each dataset at each timepoint. At 1 h, RNA-seq revealed induction of unfolded protein response pathways, consistent with early translational stress, and downregulation of differentiation/development programs (Supplementary Figure 1b). In contrast, Ribo-seq already showed enrichment of immune-related pathways (Supplementary Figure 1c), despite minimal overt immune-cell presence at this stage (Figure 1c). TE changes at 1 h highlighted pathways related to vascular function (e.g., regulation of coagulation) and transmembrane transport, alongside reduced bioenergetic and dendritic morphogenesis programs, consistent with early neuronal dysfunction (Supplementary Figure 1d). By 6 h, immune activation strengthened at both the transcriptional and translational levels (Supplementary Figure 1e-f), accompanied by marked suppression of bioenergetic/metabolic pathways (Supplementary Figure 1g) and broad perturbation of neuronal programs, cell-cycle processes, and pattern-recognition responses (Supplementary Figure 1e-g). At 24 h, immune signatures became dominant at both RNA and Ribo levels and neuronal pathway repression was more pronounced (Supplementary Figure 2a-b). In parallel, TE downregulation of translation-related pathways intensified, consistent with global translational impairment at later stages (Supplementary Figure 2c).

In summary, stroke induces a pronounced early mismatch between transcriptional and translational responses, with partial convergence by 24 h. Notably, immune pathway enrichment precedes overt neuronal loss and microglial accumulation, suggesting early activation of cell-intrinsic immune programs in response to acute ischemic stress.

### ➢ Stroke induces early synonymous codon usage changes and progressive ribosome pausing

To characterize translational dynamics after stroke, we quantified ribosome A-site pausing. Pausing increased progressively from 6 h onward and affected nearly all codons, consistent with escalating translational stress (Figure 2a-c).We next assessed codon optimality and synonymous codon usage using two complementary metrics: GC3 scores and isoacceptors codon frequencies(39). GC3, which captures third-position nucleotide bias (G/C versus A/T(39)) differed between up– and down-regulated genes only at 1 h after stroke and only in the Ribo-seq dataset (Figure 2d-f). Specifically, codon optimality showed a transient shift toward G/C-ending codons at 1 h, suggesting an early codon-selective translation program. Consistently, isoacceptors codon frequency analysis (synonymous codon usage) identified significant codon-biased translation only at 1 h and only at the translational level (Ribo-seq), with no comparable signal in RNA-seq or TE (Figure 2g).

**Figure 2:**
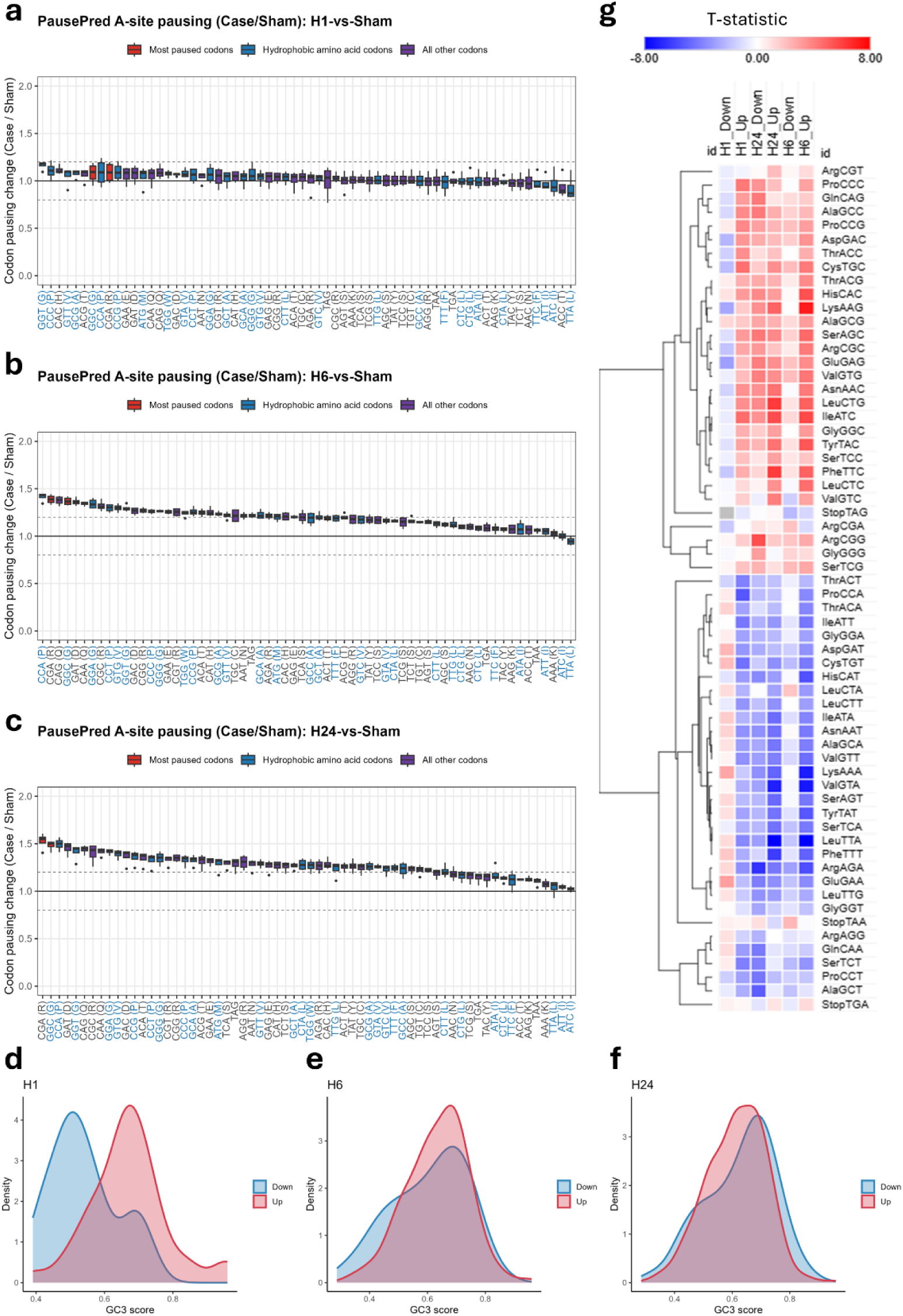
I**s**chemic **stroke leads to altered codon decoding. a-c:** Box plots showing A-site ribosome stalling after dMCAO model at each time point compared to Sham. **d-f:** Density plot of GC3 scores of the up and downregulated genes in the Ribo-seq dataset across time points (1 hour, 6 hours, and 24 hours respectively). **g:** Isoacceptors codon frequencies analysis of the up and downregulated genes in the Ribo-seq dataset across time points. T-statistics indicates significance compared to the genome average codon usage (|T-stat| ≥ 2 indicates *p* ≤ 0.05).

Together, these data indicate that stroke elicits a brief hyperacute phase of codon-biased translation, potentially reflecting an acute oxidative/metabolic stress response, followed by a later phase dominated by widespread ribosome stalling that coincides with progressive disruption of translational efficiency, including reduced TE of translation-related pathways.

### ➢ Stroke induces changes in ribosomal reading frames

Altered ribosomal reading frames can introduce premature termination codons (PTCs) and trigger nonsense-mediated decay (NMD) (Figure 3a) or generate alternative proteoforms(40–42). Ribosomal frameshifting has been linked to neurodegeneration(43), but has not been described in stroke. Given the marked perturbations in translation dynamics after stroke, we asked whether reading-frame usage changes over time. We assigned each RPF to a coding frame per gene in each sample, then computed a frameshift score (FS) reflecting deviation from the canonical 0 frame (see Methods). Using Limma-trend to compare each condition to sham, we detected a progressive increase in genes with significant shifts in dominant frame usage, emerging primarily at 6 and 24 h after stroke (Figure 3b–d, Supplementary Figure 3a). FS changes were bidirectional: some genes became more out-of-frame relative to sham (FS “UP”), whereas others became more in-frame (FS “DOWN”). We therefore focused subsequent analyses on the 6 h and 24 h timepoints. Genes with increased frameshifting were enriched for synaptic and energy metabolism pathways (Supplementary Figure 3b–c), whereas genes with decreased frameshifting were enriched for cell–cell adhesion/communication, protein localization, mitochondrial membrane organization and synaptic assembly (Supplementary Figure 3d–e). Codon-usage comparisons between FS UP and FS DOWN genes showed no major differences (Supplementary Figure 3f–h), although frameshifted genes were skewed toward specific G/C-ending codons relative to background (Supplementary Figure 3h).

**Figure 3:**
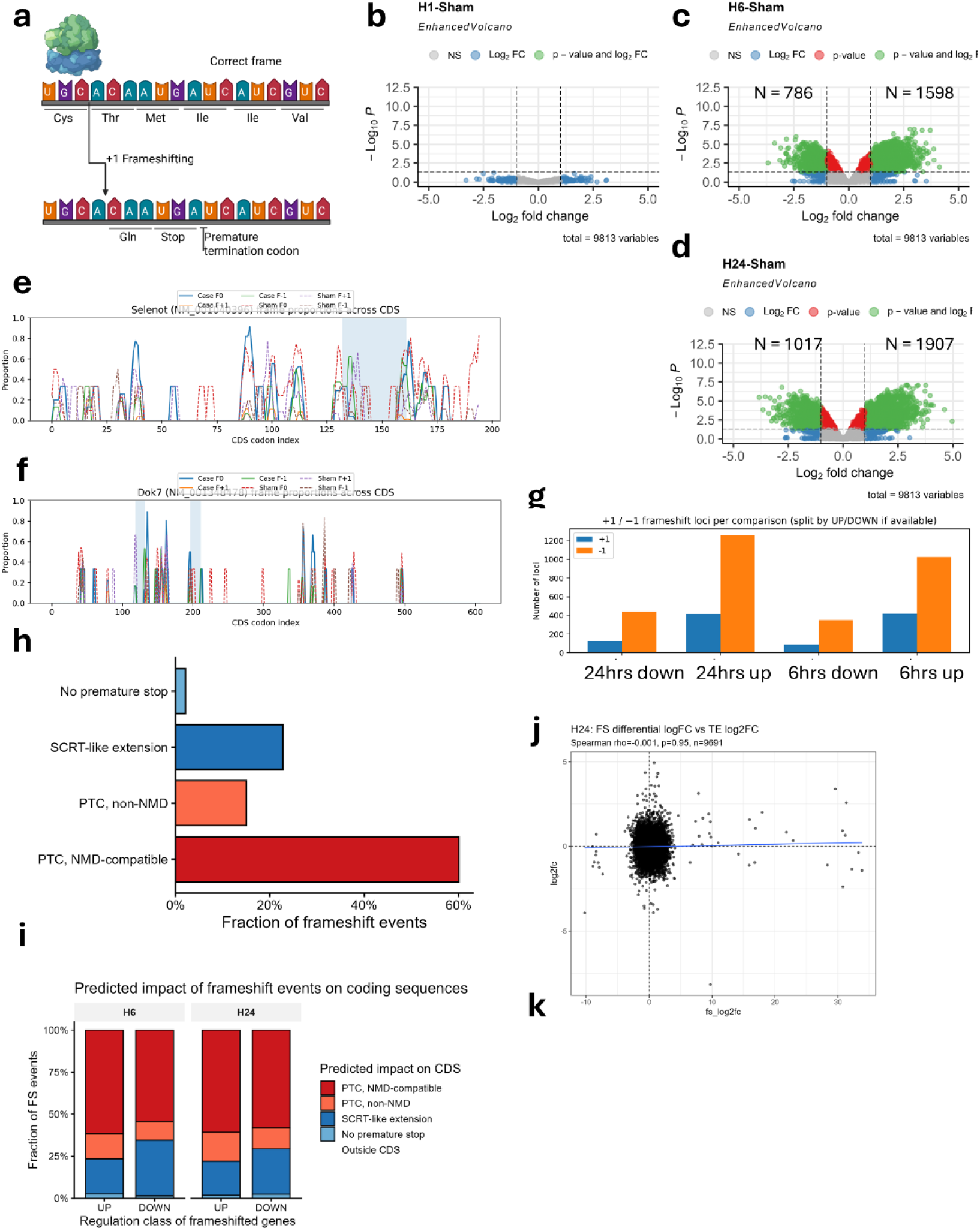
I**s**chemic **stroke leads to frameshifting events across the coding sequences. a:** Schematic showing the influence of frameshifting on the CDS reading frame. **b-d:** Volcano plots showing the changes in gene frameshifting at each time point after stroke compared to sham. Upregulated genes indicate more frameshifting (i.e. more RPFs in +1 and/or +2 frames compared to 0 frame) and vice versa. Significance was designated at |log2FC| ≥ 1 and FDR ≤ 0.05. **e:** Identification of single frameshift locus in Selenot gene, which was upregulated in the FS analysis after 24 hours. **f:** Identification of multiple frameshifting loci in Dok7 gene, which was upregulated in the FS analysis after 24 hours. **g:** Bar plot showing the number of identified +1 and –1 frameshifting events at each time point. **h:** Bar plot showing the potential impact of frameshifting on mRNA translation. This bar plot summarizes all frameshifted genes across all time points and all directions. **i**: Frameshifting impact on mRNA translation at each time point and direction of change. **j**: Spearman’s rank correlation between gene log2FC of frameshifting analysis and TE analysis at 24 hours timepoint.

To localize where frame changes occurred, we scanned the CDS of significantly frameshifted genes using an unsupervised hidden Markov model (HMM; see methods) and identified approximate windows where the dominant frame differed between stroke and sham (see Methods) (Figure 3e–f). At 6 h, we detected 228 loci in FS UP genes and 338 loci in FS DOWN genes: at 24 h, 232 and 420 loci, respectively. Some transcripts contained multiple loci. We were able to identify loci in majority of frameshifted genes (6hrs up: 1451 out of 1598 FS genes; 6hrs down: 438/786, 24hrs up: 1689/1907, and 24hrs down: 569/1017). Notably, we had lower success rate in assigning FS loci to the downregulated genes in our FS analysis than in the upregulated genes, suggesting either diffuse frame disruption that is not captured by the HMM or, alternatively, changes driven by differential ORF usage. Across timepoints and directions. −1 frame shifts were most frequently assigned than +1 frameshifting events (Figure 3g).

Using the HMM-defined windows, we modeled the impact of a frame shift on downstream coding output, focusing on PTC formation, predicted NMD susceptibility (PTC >50–55 nt upstream of the last exon–exon junction(44)), and the potential for SCRT-like C-terminal extensions via altered frame usage near stop codons. Across events, the dominant predicted outcome was PTC formation with inferred NMD susceptibility, followed by SCRT-like extension (Figure 3h–i). These results match canonical expectations for frameshifting and highlight the heterogeneity of its predicted consequences at transcriptome scale.

### ➢ Frameshifting is largely decoupled from RNA/Ribo changes

To test whether FS changes tracked with RNA abundance or ribosome occupancy, we correlated FS log2FC values with log2FC from RNA-seq, Ribo-seq and TE analyses. FS changes showed little to no correlation with gene expression, translation, or TE (Figure 3j, Supplementary Figure 4a), and overlap between FS-significant genes and conventional differential gene lists was limited (Supplementary Figure 4b–c). We next asked whether specific predicted fates (PTC, NMD, etc.) were associated with systematic shifts in RNA/Ribo metrics. K–S tests comparing log2FC distributions of fate-stratified genes to non-frameshifted genes identified modest, time– and dataset-specific effects (Supplementary Figure 4d). For example, NMD-prone events were associated with lower RNA-seq log2FC at 6 and 24 h, whereas at 6 h they were associated with higher TE log2FC. Overall, these effects were subtle and typically below thresholds for differential expression/translation, indicating that frameshifting impact cannot be inferred from RNA-seq/Ribo-seq summaries alone.

### ➢ Local coding-sequence context predicts frameshift-prone loci

To determine whether localized frameshifting events occur in reproducible sequence contexts, we trained supervised classifiers using features extracted from FS-centered coding windows. Features included codon usage, amino-acid composition, nucleotide composition, GC/GC3 content, CpG metrics, entropy, local RNA-structure metrics, motif features, and CDS-relative position. Stop-derived predictors were excluded from the final models to avoid conflating determinants of frameshifting with predicted termination consequences after frame displacement. Background windows were sampled primarily from matched CDS regions within the same transcript while excluding windows overlapping FS loci, thereby controlling for transcript-specific sequence composition.

To minimize overfitting, all models were evaluated using repeated gene-grouped 80/20 train/test splits, such that windows from the same gene were assigned exclusively to either training or held-out testing. We first compared elastic net, random forest, and XGBoost classifiers across five tasks: all FS loci versus background, FS-UP loci versus background, FS-DOWN loci versus background, FS-UP versus FS-DOWN loci, and +1 versus −1 FS loci. All three model classes performed similarly for the main FS-versus-background tasks, indicating that the signal was not specific to a single nonlinear model [Figure 4a; supplementary figure 5a]. We therefore used repeated XGBoost models for final interpretation.

**Figure 4.**
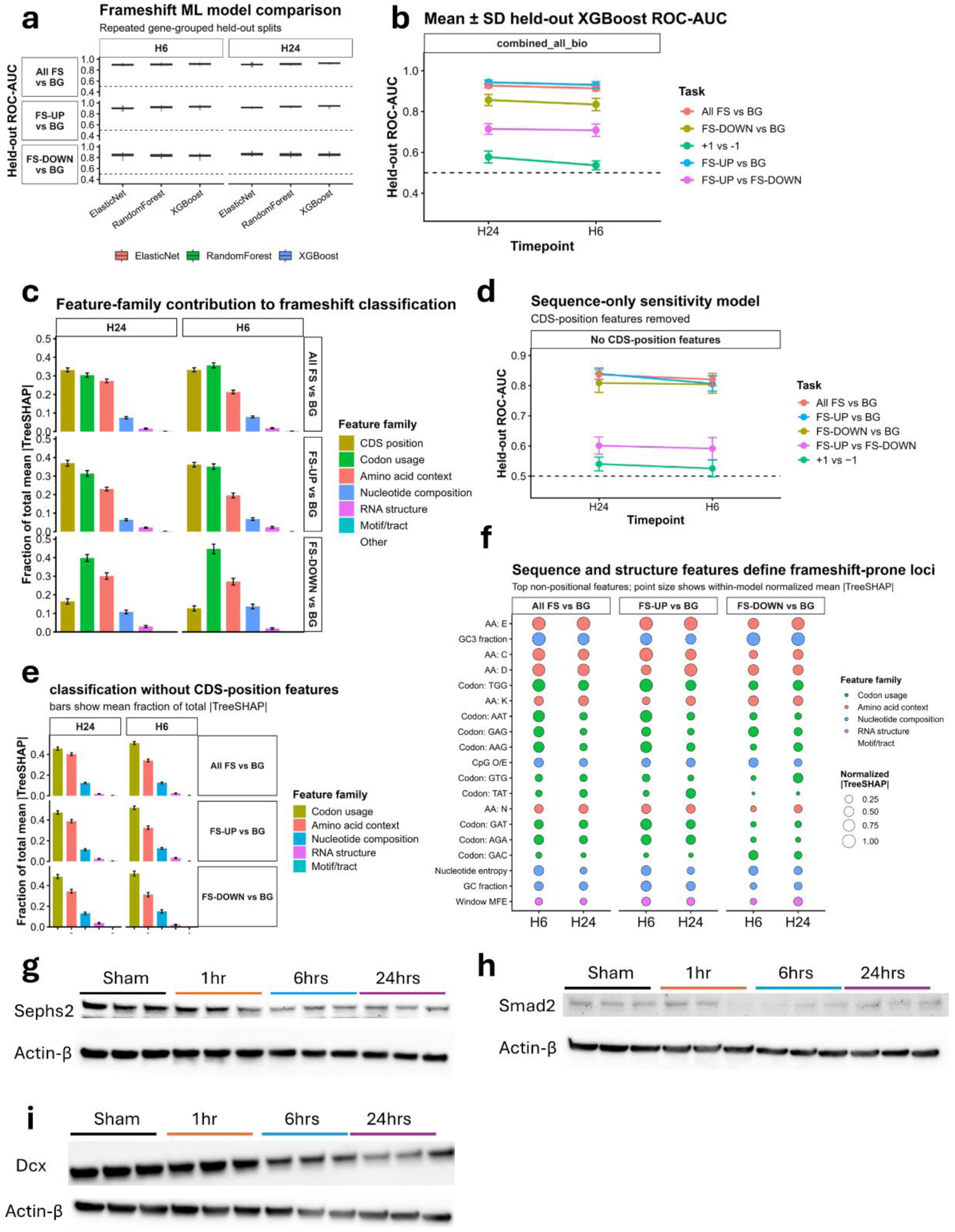
Local coding-sequence context predicts frameshift-prone loci after ischemic stroke. a: Held-out performance of elastic net, random forest, and XGBoost classifiers across repeated gene-grouped train/test splits for the indicated frameshift-classification tasks at 6 h and 24 h after dMCAO. Windows from the same gene were kept exclusively in either the training or held-out test set. **b:** Mean ± SD held-out ROC-AUC from repeated XGBoost models using the primary feature panel and same-transcript CDS-matched background windows. Dashed line indicates chance-level performance. **c:** Feature-family TreeSHAP contribution in the primary XGBoost models, showing the fraction of total mean absolute TreeSHAP signal assigned to CDS position, codon usage, amino-acid context, nucleotide composition, RNA-structure, and motif/tract features. **d:** Held-out ROC-AUC from repeated XGBoost models after excluding CDS-position features, demonstrating that local sequence and structure features retain predictive information independent of CDS-relative position. **e–f:** TreeSHAP summaries from the no-position models highlighting the dominant non-positional predictors of FS-prone loci, including amino-acid composition, codon usage, GC/GC3 content, CpG metrics, nucleotide entropy, and RNA-structure features. Point size or bar length indicates normalized or mean absolute TreeSHAP contribution across repeated held-out splits, as indicated. **g–i:** Western blot analysis and quantification of selected significantly frameshifted genes across the stroke time course: Sephs2 (g), Smad2 (h), and Dcx (i). Protein abundance was normalized to Actin-β. Protein abundance was normalized to Actin-β. Statistical analysis was performed by one-way ANOVA followed by Tukey’s post-hoc test; *P < 0.05 (See supplementary figure 5).

The strongest classification performance was observed for FS-UP versus background and all FS loci versus background, with held-out ROC-AUC values consistently above chance at both 6 h and 24 h. FS-DOWN loci were also distinguishable from background, with AUC > 0.8, although with weaker performance than FS-UP loci. In contrast, FS-UP versus FS-DOWN showed only moderate discrimination (AUC ≈ 0.7), and +1 versus −1 frame direction was predicted only weakly (AUC < 0.6) [Figure 4b]. Label-permutation models collapsed to near-chance performance, confirming that the observed classification was not explained by generic model overfitting or grouped-split structure [Supplementary figure 5b]. We focused our analysis on the strong models; All FS vs background, FS up vs background, and FS down vs background.

Held-out TreeSHAP analysis showed that FS classification was driven primarily by CDS-relative position, codon usage, amino-acid context, and nucleotide composition, with smaller contributions from RNA-structure features [Figure 4c]. In the primary same-transcript background model, CDS-relative position was a dominant predictor, particularly for FS-UP versus background and FS-UP versus FS-DOWN classification. However, models trained after excluding all CDS-position features retained strong performance in the FS all vs background, FS up vs background, and FS down vs background tasks [Figure 4d], demonstrating that local sequence composition independently distinguishes FS-prone loci. In this no-position setting, amino-acid composition, codon usage, GC3/GC content, CpG metrics, nucleotide entropy, and RNA-structure features contributed most strongly [Figure 4e-f]. Importantly, specific amino acids and codons were demonstrated to be robust predictors of frameshifted genes, indicating the contribution of local sequence/codon feature in frameshifting context. These results indicate that stroke-associated FS loci are not random CDS windows but are embedded in reproducible local coding-sequence environments.

The different classification tasks further suggested that FS-prone sequence architecture and FS direction are separable properties. Features derived from coding sequence composition robustly distinguished FS loci from matched background, especially for stroke-increased FS loci. By contrast, the weak +1 versus −1 and up vs down classifications imply that frame direction is not well explained by the current sequence-context features alone and may depend on finer event-level properties such as ribosome phase, local read distribution, or precise P-site/A-site geometry.

To test whether frameshifting corresponded to steady-state changes in protein abundance, we examined selected significantly frameshifted genes by western blotting across the stroke time course. Sephs2, which showed increased frameshifting after stroke, exhibited a modest reduction in protein abundance at 6 h and 24 h [Figure 4g; supplementary figure 5c], despite no significant change in RNA-seq, Ribo-seq, or translational efficiency [Supplementary figure 5d]. Smad2 showed increased frameshifting but no major protein abundance change [Figure 4h; supplementary figure 5e], although Ribo-seq and translational efficiency were modestly reduced at 24 h [Supplementary figure 5f]. Dcx protein levels were markedly reduced at 24 hours [Figure 4i; supplementary figure 5g], coinciding with increased frameshifting and decreased RNA abundance [Supplementary figure 5h]. Thus, individual examples did not support a simple one-to-one relationship between localized frameshifting and steady-state protein abundance.

Together, these analyses show that ischemic stroke induces reproducible, locus-resolved perturbations in ribosomal frame usage. Frameshift-prone loci are associated with local coding-sequence features and CDS positional context, but the direction and proteomic consequences of individual events are not fully captured by conventional RNA abundance, ribosome occupancy, or translational-efficiency measurements. These findings support the need for locus– and ORF-aware analyses to resolve translation-derived proteome remodeling after stroke.

### ➢ Stroke induces hyperacute stop codon readthrough events caused by termination failure

Because frameshifting can generate SCRT-like C-terminal extensions (Figure 3h), we next asked whether stroke induces stop codon readthrough (SCRT). SCRT is a well-documented but comparatively understudied phenomenon(45). It has been most extensively characterized in simpler organisms(46,47), although several functional mammalian examples exist(48–50). Despite this, large-scale efforts to quantify SCRT dynamics in multicellular physiology or disease remain limited. Here, we quantified SCRT after stroke using a custom pipeline that identifies 3′UTR RPFs and models 3′UTR-to-CDS ribosome occupancy using a GLM–beta-binomial framework to detect differential SCRT relative to sham controls (Figure 5a). Metagene analysis revealed clear 3-nt periodic P-site signal in the 3′UTR, most prominently 1 h after dMCAO (Figure 5b).

**Figure 5:**
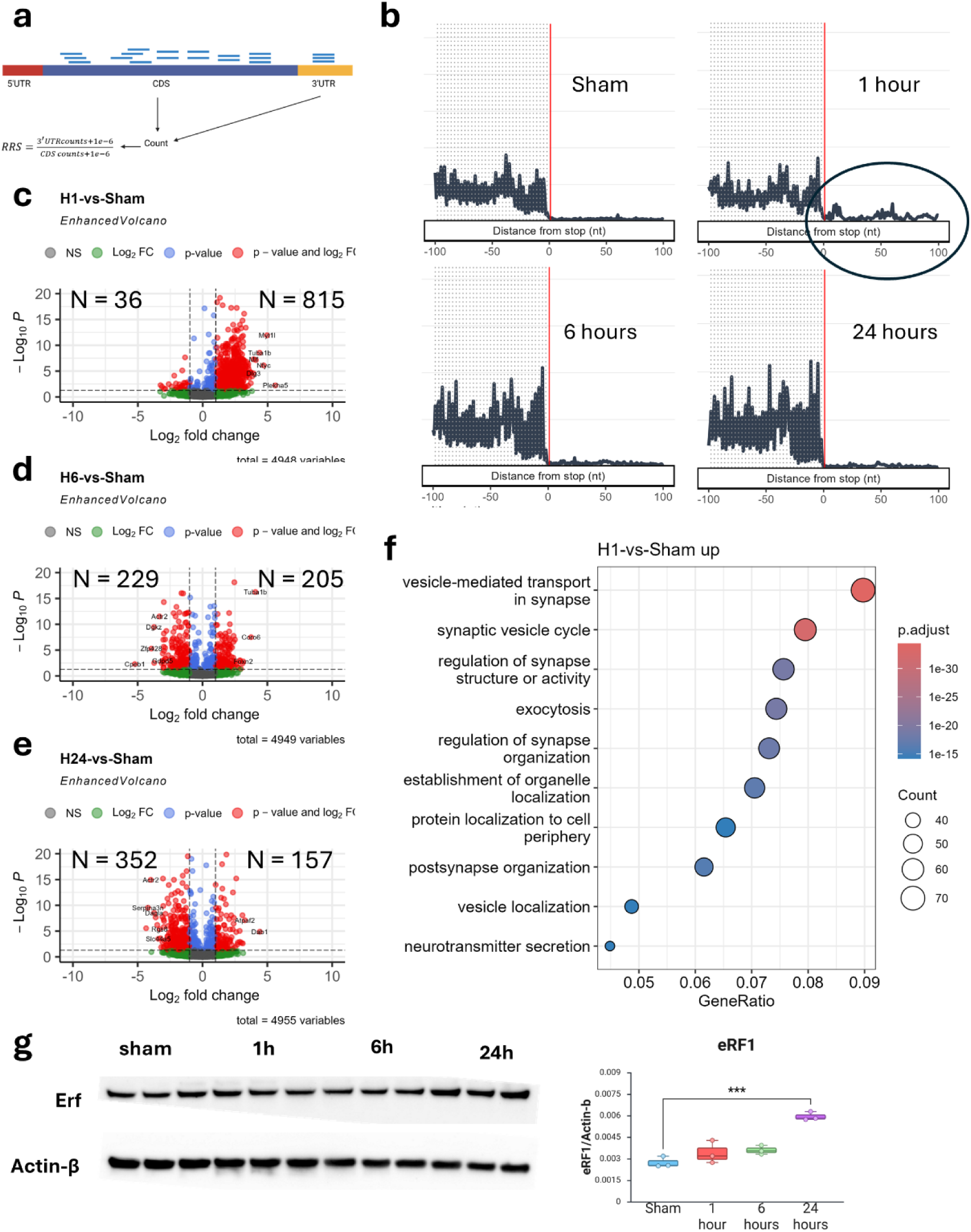
I**s**chemic **stroke leads to early burst of strop codon readthrough events. a:** Schematic representation of the approach used to identify SCRT events. **b:** Representative metagene plots showing 3-nucleotide periodicity in the 3’UTR especially in the 1-hour group. **c-d:** Volcano plots of gene showing significant SCRT events. Upregulated genes indicate more SCRT events compared to Sham and vice versa. Statistical significance cut off was |log2FC| ≥ 1 and FDR ≤ 0.05. **f:** ORA GOBP pathway analysis of upregulated genes in the SCRT analysis 1-hour after stroke. **g:** Western blot showing eukaryotic release factor (Erf or Etf1) protein levels. Protein levels remained unchanged after stroke.

Differential SCRT analysis uncovered a striking hyperacute burst at 1 h, with 815 genes showing increased SCRT relative to sham and only 36 showing decreased SCRT (Figure 5c). From 6 h onward, SCRT changes were more balanced, with a shift toward predominately decreased SCRT by 24 h (Figure 5d–e). Pathway enrichment indicated that genes with increased SCRT at 1 h were enriched for synaptic and neuronal functions (Figure 5f). At 6 h, genes with reduced SCRT were enriched for synaptic pathways, whereas genes with increased SCRT were enriched for metal ion transport, axon myelination, and metabolic processes (Supplementary Figure 6a–b). By 24 h, reduced SCRT was associated with higher cognition and synaptic functions, whereas increased SCRT was enriched for synaptic activity and nervous system development pathways (Supplementary Figure 6c–d). Together, these results argue that stroke-induced SCRT is pathway-structured rather than random, consistent with a regulated contribution to post-stroke proteome output.

### ➢ SCRT is largely independent of CDS frameshifting and reflects hyperacute termination stress

We next examined the mechanisms underlying SCRT, particularly at 1 h when the signal peaked. Because frame changes near the stop codon could produce apparent readthrough, we first assessed reading-frame usage patterns in 3′UTRs. Genes with significant SCRT showed subtle shifts in 3′UTR frame distributions after stroke (Supplementary Figure 6e). However, across samples, SCRT rate (Calculated as Ribosome Readthrough Score; RRS) and CDS frameshifting (FS) were uncorrelated (Supplementary Figure 6f), indicating that gene-level CDS frameshifting is not the primary driver of SCRT after stroke, despite the fact that some localized FS events can yield SCRT-like consequences (Figure 3h). Consistent with this, the major SCRT burst occurred at 1 h, before robust CDS frameshifting is observed, supporting the view that these are distinct (though potentially overlapping) phenomena in stroke.

We also evaluated whether changes in the canonical release factor abundance could explain the hyperacute SCRT surge. eRF1 protein levels did not show major changes early after stroke, with an observable upregulation 24 hours after stroke induction (Figure 5g), arguing against release-factor depletion as an explanation for the observed SCRT burst at 1 hour. More broadly, genes with increased SCRT at 1 h displayed a sharp pre-stop ribosome peak compared to a matched-coverage background set (Supplementary Figure 7a), a pattern absent in sham (Supplementary Figure 7b). This pre-stop aggregation was not a dominant feature at later time points (Supplementary Figure 7c–f, consistent with a transition from hyperacute, stress-driven termination failure to later, more balanced and potentially regulated readthrough dynamics.

### ➢ Sequence context and local 3′UTR structure shape SCRT dynamics

To explore how sequence and structure influence SCRT, we first compared synonymous codon usage in SCRT-significant genes to genome-wide background using Welch’s t-test. The strongest codon usage bias occurred at 1 h, where genes with increased SCRT preferentially used A/T-ending codons (Figure 6a). Codon usage bias persisted at 6 h and 24 h, but the dominant signal shifted: genes with decreased SCRT overused G/C-ending codons (Figure 6a). Because 3′UTR structure downstream of the stop codon can modulate readthrough(51), we evaluated local RNA secondary structure in the first 100 nt of the 3′UTR using RNAfold. At 1 h, genes with increased SCRT showed significantly different 3′UTR minimum free energy (MFE) relative to background, whereas genes with decreased SCRT did not (Figure 6b–c). These structure differences were not observed at 6 h or 24 h (Supplementary Figure 8a–b), again suggesting mechanistically distinct early versus late phases of SCRT regulation after stroke.

**Figure 6:**
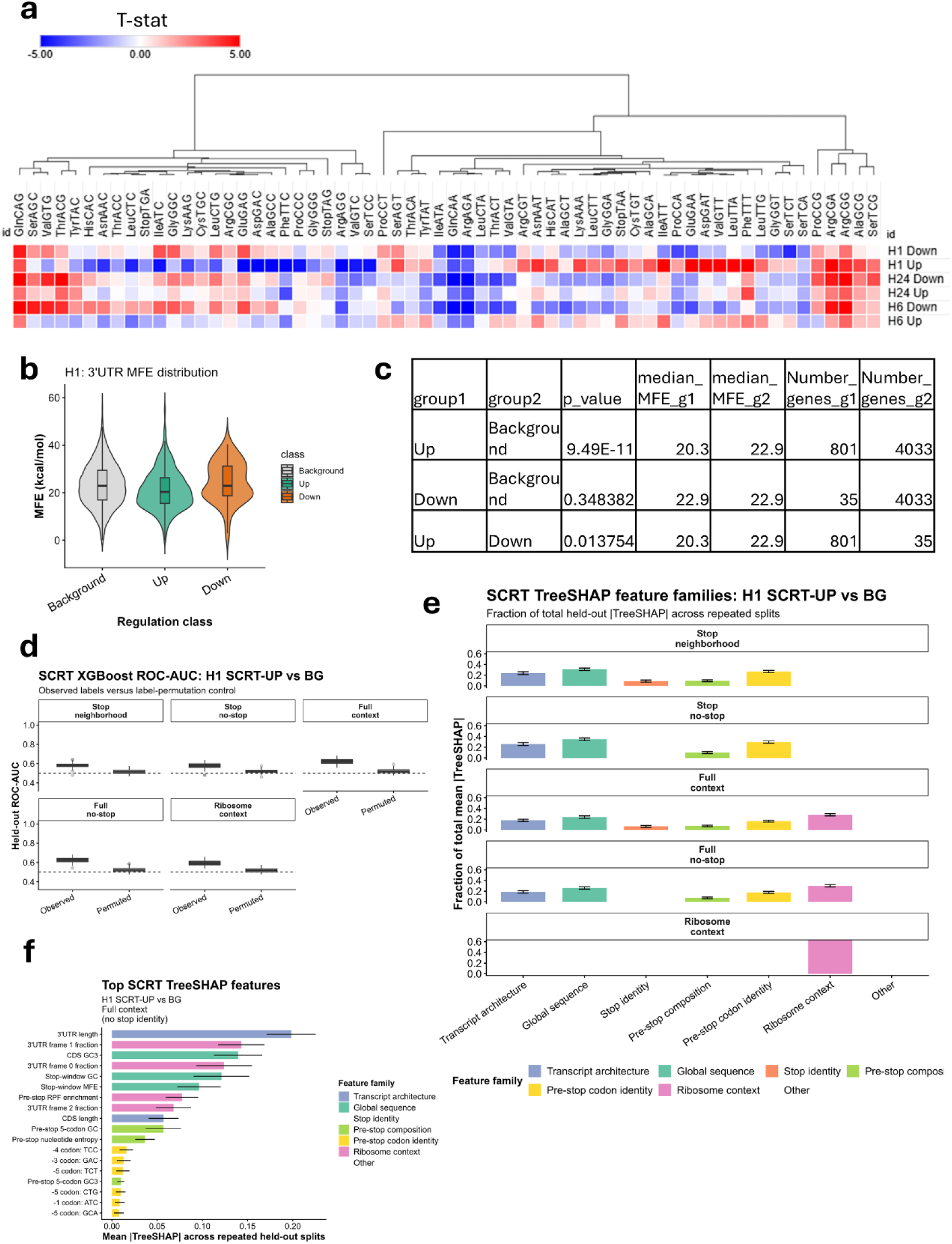
T**r**anslational **dynamics and 3’UTR structures predict SCRT potential. a:** Isoacceptors codon frequencies analysis of genes with significant SCRT events. **b:** Violin plot of minimum free energy (MFE) analysis of RNAfold using the first 100 nucleotides of the 3’UTR as input. **c:** Wilcoxon test analysis of MFE comparing the up and downregulated genes to the background or to each other. **d:** XGBoost held-out ROC-AUC for classifying H1 SCRT-UP genes against coverage-matched background genes across five feature panels. Observed-label models are compared with label-permutation controls using repeated gene-grouped held-out splits. **e:** TreeSHAP feature-family contribution for the H1 SCRT-UP versus background task across feature panels. Bars show the fraction of total mean absolute TreeSHAP signal assigned to each feature family. **f:** Top TreeSHAP features from the primary H1 SCRT-UP versus background model using the full-context feature panel with stop codon identity removed. Error bars indicate variation across repeated held-out splits.

### ➢ Machine learning identifies multifactorial transcript-context determinants of SCRT

Given the number of potentially interacting features associated with SCRT, we next applied a machine-learning framework to identify transcript features that distinguish SCRT-responsive genes from matched background genes. We first compared elastic-net, random forest, and XGBoost classifiers across timepoints and SCRT classification tasks. XGBoost showed competitive held-out performance and was selected for the final analysis because it enabled consistent TreeSHAP-based interpretation across repeated gene-grouped train/test splits (Supplementary Figure 8c). We then trained final XGBoost models using five feature panels: stop-neighborhood sequence, stop-neighborhood sequence without stop codon identity, full transcript context, full transcript context without stop codon identity, and ribosome-context-only features. Final models were evaluated across 100 repeated gene-grouped iterations, and model performance and feature importance were summarized as the mean ± SD across iterations.

The strongest and most reproducible SCRT prediction was observed at 1 h after stroke, particularly for the SCRT-UP versus matched-background task (Figure 6d; Supplementary Figure 8d). Full-context and stop-neighborhood models performed above label-permutation controls, and this separation was retained after removing stop codon identity. Thus, the classifier was not simply learning canonical TAA/TAG/TGA usage. Ribosome-context-only models also carried predictive signal but were generally weaker than models incorporating transcript architecture and sequence-context features, indicating that 3′UTR frame usage and pre-stop ribosome enrichment contribute to SCRT susceptibility but do not fully explain it alone (Figure 6d). Across 6h and 24h, model performance was weaker and more heterogeneous than in the 1h SCRT-UP susceptibility model (Supplementary Figure 8d). However, SCRT-UP versus SCRT-DOWN models at H24 showed moderate separation, suggesting that late SCRT directionality still retains transcript-architecture and sequence-context structure.

TreeSHAP analysis showed that SCRT prediction was distributed across several feature families rather than dominated by a single variable class (Figure 6e). In the primary 1h SCRT-UP versus background full-context model without stop codon identity, the top features included 3′UTR length, 3′UTR frame usage, CDS GC3, stop-window GC content, stop-window predicted structure, pre-stop ribosome enrichment, CDS length, and pre-stop nucleotide/codon composition (Figure 6f). Inclusion of stop codon identity introduced stop-codon features among the interpretable variables, but did not eliminate the contribution of transcript architecture, global sequence composition, ribosome context, and pre-stop codon features (Supplementary Figure 8e). In the 1h SCRT-UP versus SCRT-DOWN model, 3′UTR length, CDS GC3, and stop-proximal RNA structure contributed to class separation despite the weaker overall model performance (Supplementary Figure 8f). At 24h, the SCRT-UP versus SCRT-DOWN model highlighted CDS length, CDS GC3, and pre-stop nucleotide entropy among the main discriminating features (Supplementary Figure 8g). Collectively, these results indicate that SCRT susceptibility is associated with a multifactorial transcript-context signature involving sequence composition, local RNA structure, transcript architecture, and dynamic ribosome features. The modest overall model performance also indicates that additional determinants of SCRT induction remain to be identified.

Finally, SCRT changes at 1 h showed no significant correlation with RNA-seq or Ribo-seq differential signals (Supplementary Figure 8h–j), indicating that the hyperacute SCRT burst is largely orthogonal to conventional expression/translation changes. At 6 h and 24 h, the strongest association was a modest negative correlation between SCRT and Ribo-seq log2FC (ρ = −0.289 and −0.282, respectively; p < 0.0001; not shown). To assess functional coupling to downstream regulation, we compared RNA-seq, Ribo-seq, and TE log2FC distributions for SCRT-altered genes using Wilcoxon rank-sum and Kolmogorov–Smirnov tests with Benjamini–Hochberg correction (Supplementary Figure 9). SCRT changes had no detectable effect on RNA abundance (Supplementary Figure 9a) but were associated with changes in translation (Supplementary Figure 9b) beginning at 6 h and in TE (Supplementary Figure 9c) beginning at 1 h. Increased SCRT tended to reduce TE, whereas decreased SCRT tended to increase TE, indicating that SCRT-associated effects are subtle yet remain primarily translational.

### ➢ Stroke induces changes in ORF dominance and translation

We next examined how stroke alters open reading frame (ORF) usage and translation. Our gene-level frameshifting analysis indicated that although thousands of genes showed altered frame usage, discrete frameshift loci could be localized in only a subset, suggesting that much of the signal may reflect altered ORF usage rather than ribosomal slippage. We therefore used RiboCode(36) to identify translated ORFs in each sample. ORFs were filtered for length ≥10 amino acids (≥30 nt), P-sites ≥20, p ≤ 0.05, and ≥50% of reads in frame 0. Across all samples we detected ∼40,000 translated ORFs, dominated by annotated ORFs, followed by novel ORFs and downstream ORFs (dORFs) in the 3′UTR (Figure 7a). Annotated ORFs were the longest (mean 660 aa; median 504 aa) (Figure 7b). dORFs and overlap-dORFs were the next longest, whereas novel ORFs were ∼one-third the length of annotated ORFs on average (Figure 7b).

**Figure 7:**
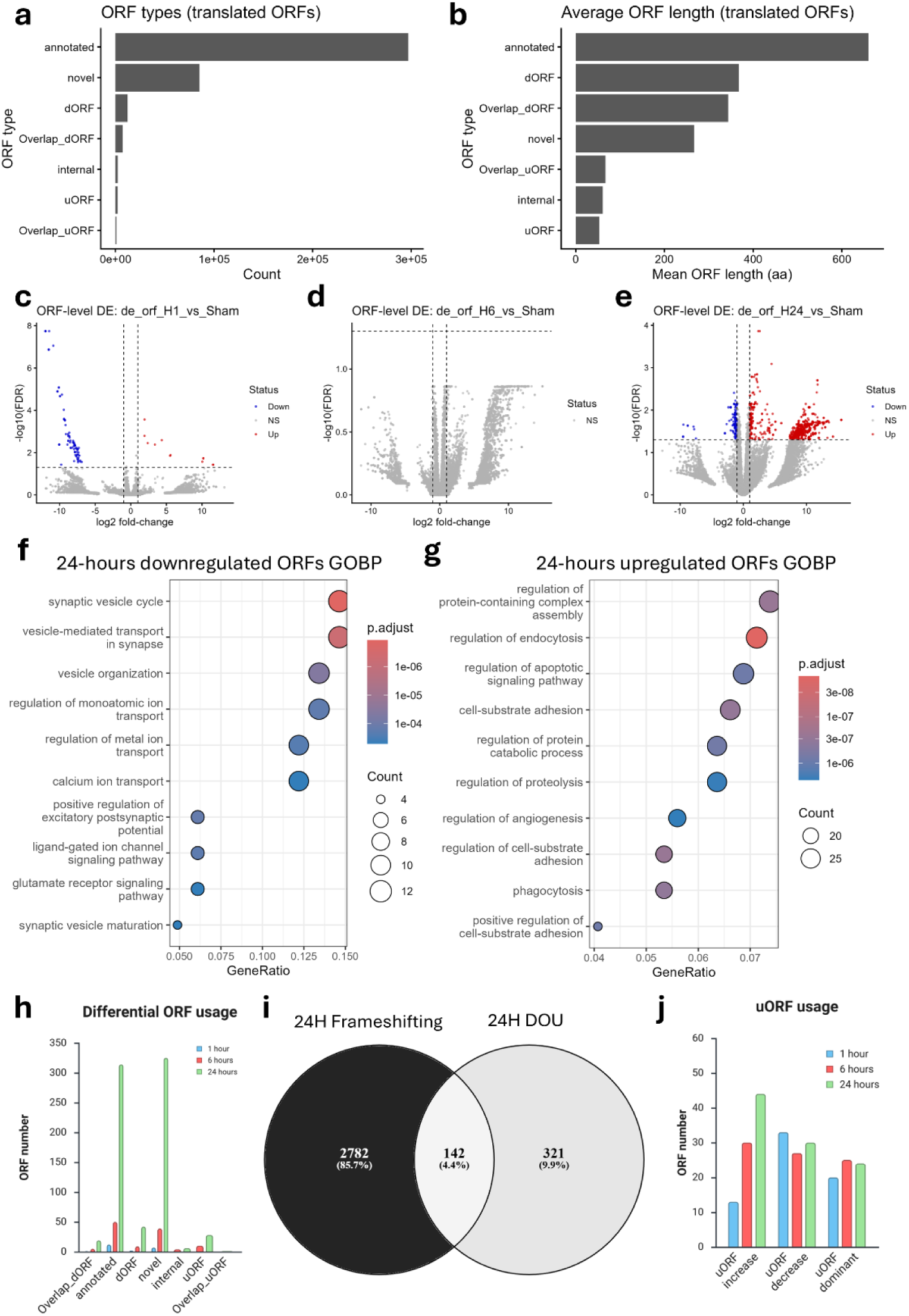
I**s**chemic **stroke leads to transcriptome-wide ORF changes. a:** Bar plot showing the ORF subtype counts detected in the Ribo-seq dataset. **b:** Bar plot showing the average translated ORF length for each subtype detected. **c-e:** Volcano plot showing the differentially translated ORFs at each timepoint compared to sham. **f-g:** ORA GOBP pathway analysis of downregulated and upregulated ORFs (using the Gene symbols as input) at the 24-hours timepoint. **h:** Bar plot showing the number of genes with differential ORF usage (DOU) detected by DRIMSeq and the subtypes of the ORFs. **i:** Venn diagram showing the overlap between genes having significant DOU and frameshifting at 24-hour timepoint. **j:** Bar plot showing genes with uORF usage changes across time points. uORF dominance is a separate entity from uORF increase and decrease, it shows genes with their dominant ORF being uORF regardless of change in usage after stroke.

We first performed an ORF-level differential analysis using total P-site counts per ORF as input. At 1 h, most significant ORFs were downregulated with few upregulated (Figure 7c). At 6 h, we detected essentially no differentially translated ORFs (Figure 7d). In contrast, 24 h showed robust ORF-level remodeling with many ORFs significantly upregulated (Figure 7e). In both the 1 h and 24 h comparisons, annotated ORFs were the most frequent differentially translated class, followed by novel ORFs and dORFs (Supplementary Figure 10a–b). Gene Ontology (GOBP) over-representation analysis of parent genes revealed that ORFs downregulated at 1 h were enriched for growth hormone signaling, focal adhesions, and cell junction pathways (Supplementary Figure 10c), and KEGG analysis highlighted axon guidance as the top pathway (Supplementary Figure 10d), linking early ORF suppression to neuronal circuitry. By 24 h, downregulated ORFs were enriched for neuronal and synaptic functions (Figure 7f), whereas upregulated ORFs were enriched for apoptotic signaling, regulation of protein metabolism, phagocytosis, and cell–substrate adhesion (Figure 7g), consistent with concurrent neuronal dysfunction/death and immune activation/clearance as injury consolidates.

### ➢ Stroke drives progressive differential ORF usage (DOU) independent of RNA abundance

Because a single gene can encode multiple translated ORFs (and multiple isoforms can expand this space), we next tested whether stroke shifts dominant ORF usage. We defined “canonical” ORFs as those accounting for ≥60% of all P-sites attributed to a gene within a sample. We then used DRIMSeq(38) to detect genes with significant shifts in canonical ORF usage. Differential ORF usage (DOU) increased progressively from 1 h and peaked at 24 h (Figure 7h). Notably, DOU increased at 6 h relative to 1 h despite the absence of ORF-level differential translation at 6 h (Figure 7c–d), indicating that ORF remodeling can occur even when total ORF translation changes are minimal.

Across DOU events, annotated and novel ORFs dominated, reflecting their prevalence in the discovery set, followed by dORFs. Unlike our SCRT analysis, switching toward dORF dominance was significantly increased at 24 h, suggesting regulated downstream initiation rather than deregulatory readthrough. To determine whether DOU tracked with conventional regulatory modes, we tested enrichment of DOU among genes with significant RNA-seq and/or Ribo-seq changes using Fisher’s exact tests with FDR correction. At 1 h and 6 h, DOU was not significantly enriched in any regulatory mode (Supplementary Figure 10e). At 24 h, DOU was modestly enriched among genes without significant RNA or Ribo changes (OR = 1.5, q = 0.017; 589 DOU genes not significant), whereas transcription-only and combined RNA+Ribo modes were depleted for DOU (OR = 0.16 and 0.1; q = 6.27E−07 and 0.015, respectively), although the absolute counts were small (5/1075 and 1/340). These results indicate that ORF usage shifts are largely orthogonal to changes in mRNA abundance or bulk ribosome occupancy. Consistent with this interpretation, ∼30% of genes with significant DOU at 24 h also scored significant in our frameshifting analysis (Figure 7i), indicating that gene-level frame changes capture a substantial component of ORF switching after stroke.

### ➢ uORF engagement increases at 24 h and associates with reduced coding translation

Within the DOU results, we observed an increase in uORF usage at 24 h. uORFs can repress downstream CDS translation(52,53). We therefore stratified genes showing increased uORF usage, decreased uORF usage, or switching to uORF dominance (Figure 7j). To relate these patterns to gene regulation, we compared RNA-seq, Ribo-seq, and TE log2FC distributions for uORF-changing genes versus stable genes using Wilcoxon rank-sum and Kolmogorov–Smirnov tests with Benjamini–Hochberg correction. The strongest association was at the translational level: genes with either increased or decreased uORF usage showed lower median Ribo-seq log2FC than stable genes beginning at 6 h (Supplementary Figure 11a). TE effects were most pronounced for uORF-increased genes, which showed reduced TE at 24 h (Supplementary Figure 11b). RNA-seq effects were weakest and only evident at 24 h (Supplementary Figure 11c). Although the number of genes switching toward uORFs was modest, these genes were enriched for synaptic functions at 24 h (not shown), supporting biological specificity.

### ➢ Gene-level upstream translation increases progressively and represses CDS translation

To complement ORF calling, we quantified upstream translation at the gene level using a GLM–beta-binomial framework analogous to SCRT, comparing 5′UTR P-sites to CDS P-sites. This analysis revealed progressive changes in upstream translation that peaked at 24 h (Figure 8a–c), mirroring the ORF-based remodeling. Clustering of the upstream translation ratio (UPR; 5′UTR/CDS reads per gene) separated late timepoints from sham and 1 h (Figure 8d). Pathway analyses highlighted enrichment for neuronal and synaptic programs (Figure 8e–f, Supplementary Figure 12a–d). We also observed overlap between genes with significant frameshifting at 24 h and genes with significant upstream translation changes (Figure 8g; significance: |log2FC| ≥ 1, FDR ≤ 0.05).

**Figure 8:**
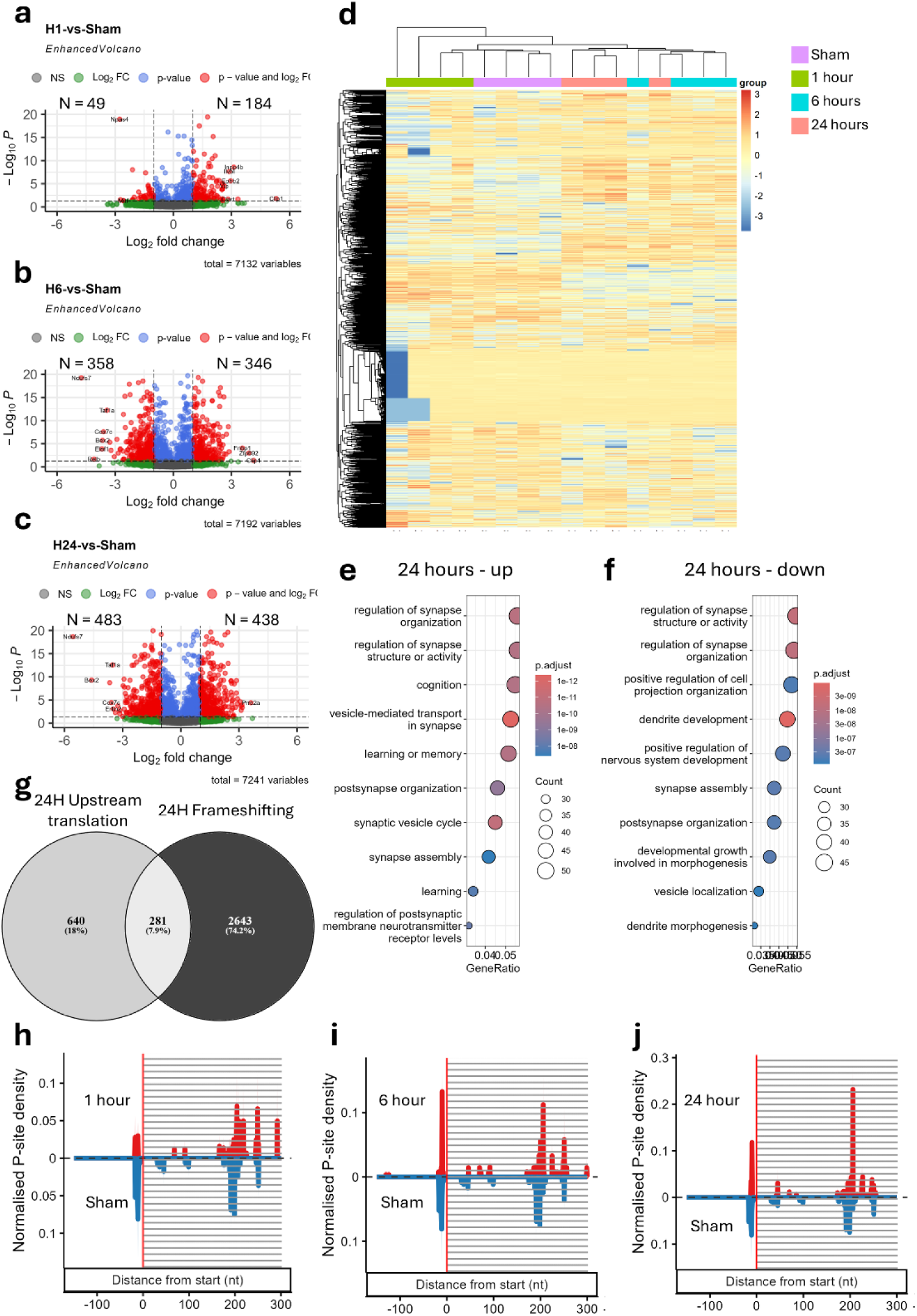
I**s**chemic **stroke leads to progressive upstream translation starts. a-c:** Volcano plots showing the number of genes with increased or decreased upstream (5’UTR) RPF ratio compared to CDS RPFs at each time points. **d:** Heatmap showing the gene upstream translation ratio (UTR; ratio of 5’UTR reads to CDS reads per gene). **e-f:** ORA GOBP analysis of up and downregulated genes in the upstream translation analysis at the 24-hour timepoint. **g:** Venn diagram showing the overlap between genes having significant frameshifting and those significant in the upstream translation analysis. **h-j:** Normalized P-side density mirror plots of DCX gene, which is significant in both frameshifting analysis and upstream translation analysis across timepoints.

To visualize locus-level examples, we examined Dcx and Smad2, both of which showed frameshift and Dcx showed reduction in protein abundance (Figure 4f–g). For Dcx, 1 h showed a reduced 5′UTR P-site peak relative to sham (Figure 8h), followed by the strongest relative increase at 6 h (Figure 8i) that persisted at 24 h (Figure 8j). Smad2 showed a similar pattern, with 5′UTR peaks comparable to sham at 1 h and increased peaks from 6 h onward (Supplementary Figure 12e–g). Finally, consistent with uORF-mediated gating, differential upstream translation (DUT) was associated with reciprocal shifts in CDS translation and TE: genes with increased DUT showed lower Ribo-seq and TE log2FC values (and vice versa), detectable from 1 h onward (Supplementary Figure 13a–b). No consistent association was observed with RNA abundance (Supplementary Figure 13c), indicating that upstream translation remodeling primarily acts at the translational level.

Collectively, these results indicate that stroke induces a time-dependent ORF remodeling program that is modest in the hyperacute phase but becomes prominent from 6–24 h. This program includes shifts in dominant ORF usage, increased uORF engagement, and rising upstream translation that inversely tracks CDS translation and TE (without parallel RNA changes), consistent with translational gating that diversifies coding outputs and contributes to proteome remodeling during stroke.

## Discussion

In this work, we provide a time-resolved analysis of translational remodeling during stroke that reveals a staged translational response program not apparent from RNA abundance analysis alone. We show that changes in mRNA expression are largely disconnected from mRNA translation during the hyperacute and acute phases after stroke. We also identify a short-lived, codon-biased translational program in the hyperacute phase, distinct from what is observed in cellular oxidative stress models(16,54). In parallel, we detect pervasive stop codon readthrough events consistent with translation termination failure, with the potential to alter C-terminal structures of neuronal and synaptic proteins before major changes in their expression. At later stages, we observe broad changes in ribosomal reading frames. A subset of frameshifting events can be localized to discrete CDS regions, consistent with ribosomal slippage triggered by stalling or collision, coinciding with progressive A-site pausing. These localized events often introduce downstream premature termination codons (PTCs) and, in some cases, predict nonsense-mediated decay. Frameshifting overlapped substantially with progressive alternative ORF usage, suggesting that alternative ORF selection rather than could explain the apparent frame changes. Finally, using two orthogonal approaches, we observe progressive increases in uORF usage and upstream translation initiation. Increased upstream translation (increased 5′UTR RPFs) is inversely associated with CDS translation and translational efficiency, consistent with previous observations(52,53). Collectively, these findings indicate that translational changes in the ischemic brain extend well beyond mRNA/protein abundance, reflecting complex regulatory and deregulatory processes that shape translational efficiency and proteome stability and diversity. To our knowledge, these phenomena have not been systematically explored in *in vivo* stroke models, and rarely in mammalian disease contexts. Thus, our analysis provides a framework for extracting hidden translational mechanisms from Ribo-seq datasets, applicable beyond stroke and neurological disorders.

Despite modern improvements in stroke therapy, stroke remains the second leading cause of morbidity and mortality worldwide(4), underscoring the need for novel neuroprotective strategies(55). A major barrier to therapeutic development is an incomplete understanding of the evolving molecular dynamics of stroke pathophysiology. Stroke is temporally dynamic and involves interactions among multiple cell types(5). Nevertheless, stroke molecular studies remain dominated by transcriptomics and proteomics(56,57), with recent shifts toward single-cell and spatial transcriptomics(58,59). While these approaches are crucial for defining cellular states, they do not resolve the mechanisms controlling mRNA translation, particularly under acute oxidative and metabolic stress(60,61), conditions central to ischemic injury. Our observation of transcription–translation discordance further emphasizes the importance of studying translation regulation directly. To date, only a few studies have examined translation in stroke(62,63), primarily through translatome capture in Ribo-tag mice(62,63). Although informative for cell-specific ribosome loading, these approaches lack nucleotide-resolution ribosome footprinting and therefore cannot capture the breadth of translational regulatory and deregulatory phenomena uncovered here. Nonetheless, similar observations were made in oxygen glucose deprivation (OGD) model using PC12 rat neuronal cells(64), further supporting the notion that our observations are not sequencing artifacts, but reproducible molecular phenomena across *n vitro* and *in vivo* stroke models.

Using ribosome profiling, we identify multiple translational behaviors that are not accessible to transcriptomics or standard proteomics. Importantly, these phenomena change dynamically over time, revealing a nuanced landscape as neurons respond to acute ischemic stress, remodel the translatome, and ultimately fail. In the hyperacute phase, we observe stress-responsive codon-biased translation with a shift toward G/C-ending codons(12,16) and a pronounced spike in stop codon readthrough. SCRT generates C-terminally extended proteoforms and has been described in viral infection, hypoxia, and oxidative stress, particularly in prokaryotes and yeast, but has not been systematically studied in multicellular organisms or in acute brain injury(64–67). Our data reveal widespread readthrough acutely after stroke, consistent with an early stress response(68) followed by a shift toward more balanced readthrough patterns at later timepoints, suggestive of additional regulatory processes. ML analysis implicated 3’UTR sequence features, G/C-ending codon usage, and pre-stop ribosome stacking and termination failure followed by frameshifting as key drivers of hyperacute SCRT(51,68,69). SCRT has been proposed as a mechanism that expands proteome diversity(70), and functional examples exist in neurons, including AQP4 and its readthrough isoform AQP4ex(49). We detect later SCRT remodeling in neuronal and synaptic genes, with both increases and decreases relative to sham. However, these later dynamics differ from the hyperacute burst, which appears dominated by termination stress. Notably, SCRT-associated genes exhibit significant changes in translational efficiency without detectable changes in mRNA expression, reinforcing that SCRT is primarily a translational phenomenon independent from mRNA expression. Whether these events yield stable proteoforms and alter gene function will require targeted proteomics and functional studies.

After the hyperacute phase, we observe progressive frameshifting beginning 6 h after stroke. A subset of events can be localized to discrete CDS loci, consistent with ribosome slippage or stalling and/or collision, coinciding with progressive A-site pausing(71). Ribosome stalling and collisions canonically trigger the ribotoxic stress response(60,72,73), which limits further ribosome loading. We did not detect overt activation of this pathway in the stroke model (data not shown), suggesting that collision signaling may be subtle, transient, or restricted to specific transcripts, consistent with the fact that localizable frameshift loci were not detected in all of the frameshifted genes. ML analyses further indicate that frameshifting is strongly shaped by local sequence features, including nucleotide and codon composition, amino-acid content, and RNA structure, and distinguish loci with increased versus decreased frameshifting relative to sham. Consequence modeling predicts frequent downstream PTC formation and potential NMD engagement. However, not all gene-level frameshifting signals could not be mapped to discrete loci, raising the possibility that frame changes could be diffuse or, more plausibly, reflective alternative ORF usage. Indeed, differential ORF usage (DOU) increases progressively with stroke, and overlaps substantially with frameshifting calls, supporting the interpretation that alternative ORF selection explains most apparent frame disruption, whereas collision/slippage-driven frameshifting represents a subset in stroke. Consistent with this remodeling, our ORF analyses reveal progressive uORF engagement. uORFs are established regulators of downstream translation(52,53). Using a GLB-beta-binomial model, we identify genes with increased 5′UTR RPF proportions relative to the CDS after stroke, enriched for neuronal and synaptic pathways. Importantly, increased or decreased upstream translation correlates inversely with translational efficiency, consistent with prior work linking upstream initiation to suppression of CDS translation(52,53,74,75).

Collectively, our data expand the molecular framework of ischemic stroke by placing translation control and translation failure mechanisms alongside transcriptional programs and inflammatory remodeling. Our results support a temporally ordered cascade of termination stress, elongation/collision stress, and ORF rewiring—translation-layer mechanisms extensively characterized in cell stress models but rarely mapped in vivo in mammalian disease. More broadly, our study highlights the need for complementary tools to validate these mechanisms in stroke and other disorders, and suggests that orthogonal biochemical, genetic, and proteomic approaches will be essential to identify regulators and establish causal links.

Several limitations should be considered. First, ribosome profiling was performed on bulk infarct-containing tissue; thus, observed changes may reflect both cell-intrinsic regulation and evolving cell-type composition, and future cell-type-resolved translatomics or spatial approaches will refine attribution. Second, while SCRT-like signals and 3′UTR ribosome occupancy are consistent with termination failure, rescue, and/or impaired recycling, ribosome footprints alone cannot distinguish productive readthrough that yields stable C-terminally extended proteoforms from rescue-associated post-termination ribosomes or scanning/reinitiation in 3′UTRs, as documented in other systems(69). Third, localized out-of-frame signatures are consistent with slippage/collision at specific loci, but we do not directly measure collisions or ribotoxic signaling; disome profiling or biochemical readouts would strengthen mechanistic specificity(61,76). Fourth, consequence annotation (e.g., PTC/NMD susceptibility) and ORF assignment depend on transcript models and general NMD rules and should be interpreted as predicted propensity rather than direct measurements; isoform-resolved transcription/decay profiling and targeted validation would refine these predictions(77). Finally, we focus on discrete timepoints in a single model, using adolescent mice, and only in male sex; expanded time courses, reperfusion paradigms, additional regions, female stroke models, using aged mice, and other stroke subtypes will be important to generalize the staged model.

In summary, time-resolved integration of RNA-seq and ribosome profiling shows that translational remodeling during ischemic stroke is not a uniform suppression of protein synthesis, but a staged program evolving from early termination/3′UTR ribosome stress and codon-biased translation to later elongation stress, out-of-frame decoding signatures, and progressive rewiring of ORF and uORF usage. These layers help explain why transcript abundance incompletely predicts translational output during acute brain injury and highlight translational control nodes, termination/recycling, collision surveillance, and ORF selection, that may shape neuronal proteostasis and fate as ischemia progresses. More broadly, our framework demonstrates how nucleotide-resolution ribosome footprinting can reveal hidden translational regulatory and deregulatory mechanisms in vivo, providing a foundation for mechanistic testing across diverse pathologies beyond stroke.

## Acknowledgments

Large language mode (ChatGPT) was used for language editing and as assistant in code development. ChatGPT was used to generate graphical abstract, which was thoroughly evaluated by the authors. The authors take full responsibility for all the content of the manuscript. The authors would like to thank Ms Natsumi Konno and Dr Michitoshi Watanabe for their assistance in the various stages of this work. This work was supported by Biomedical Research Core of Tohoku University Graduate School of Medicine.

## Authors contributions

**S.R.:** Conception and study design. Writing. Bioinformatics. Funding. Administration. Supervision. **Y.K., T.N., D.A., A.M.,** and **H.I:** Experimentation and animal modeling. **H.E.:** critically reviewed the manuscript. Supervision. **K.N.:** Critically reviewed the manuscript. Administration. Funding.

## Data availability

Raw Ribo-seq and RNA-seq fastq files were deposited in sequence read archive (SRA; PRJNA1220332 and PRJNA1402676).

## Code availability

The python scripts used to calculate frameshifting and analyze the frameshift location and impact are available through GitHub (https://github.com/Rashad-lab/Gene-level-frameshifting). All other analysis were performed using R language programming and available libraries. Further information can be provided by the corresponding author upon reasonable requests.

