## Supplementary figures for "Ischemic stroke induces staged translational remodeling in mouse cortex independent from transcriptional responses"

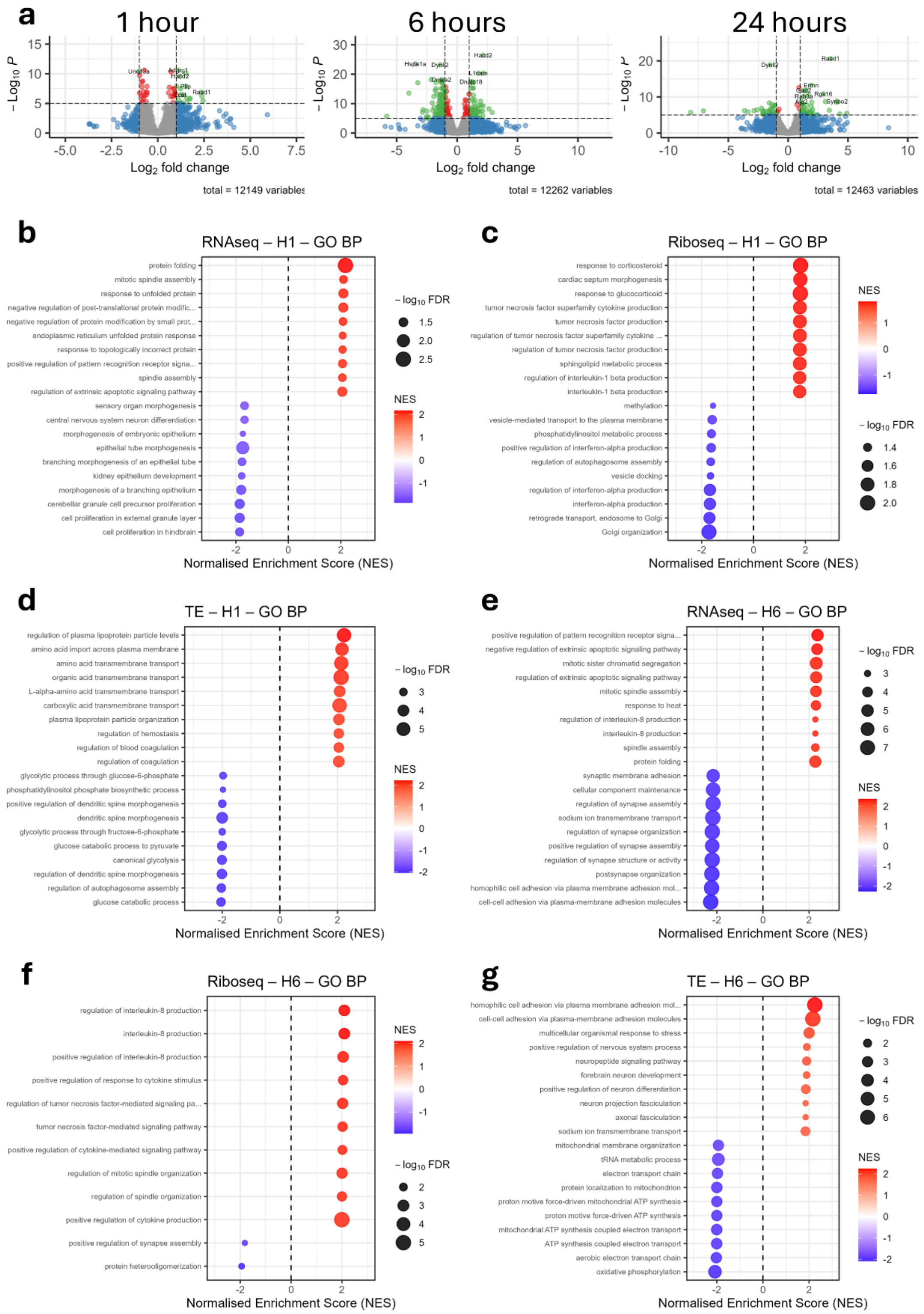


**Supplementary figure 1: a:** Volcano plots of translational efficiency changes over time after stroke induction. **b-d:** Pre-ranked GSEA analysis using GOBP gene set on different gene regulatory modes 1 hour after stroke. **e-g:** Pre-ranked GSEA analysis using GOBP gene set on different gene regulatory modes 6 hours after stroke.

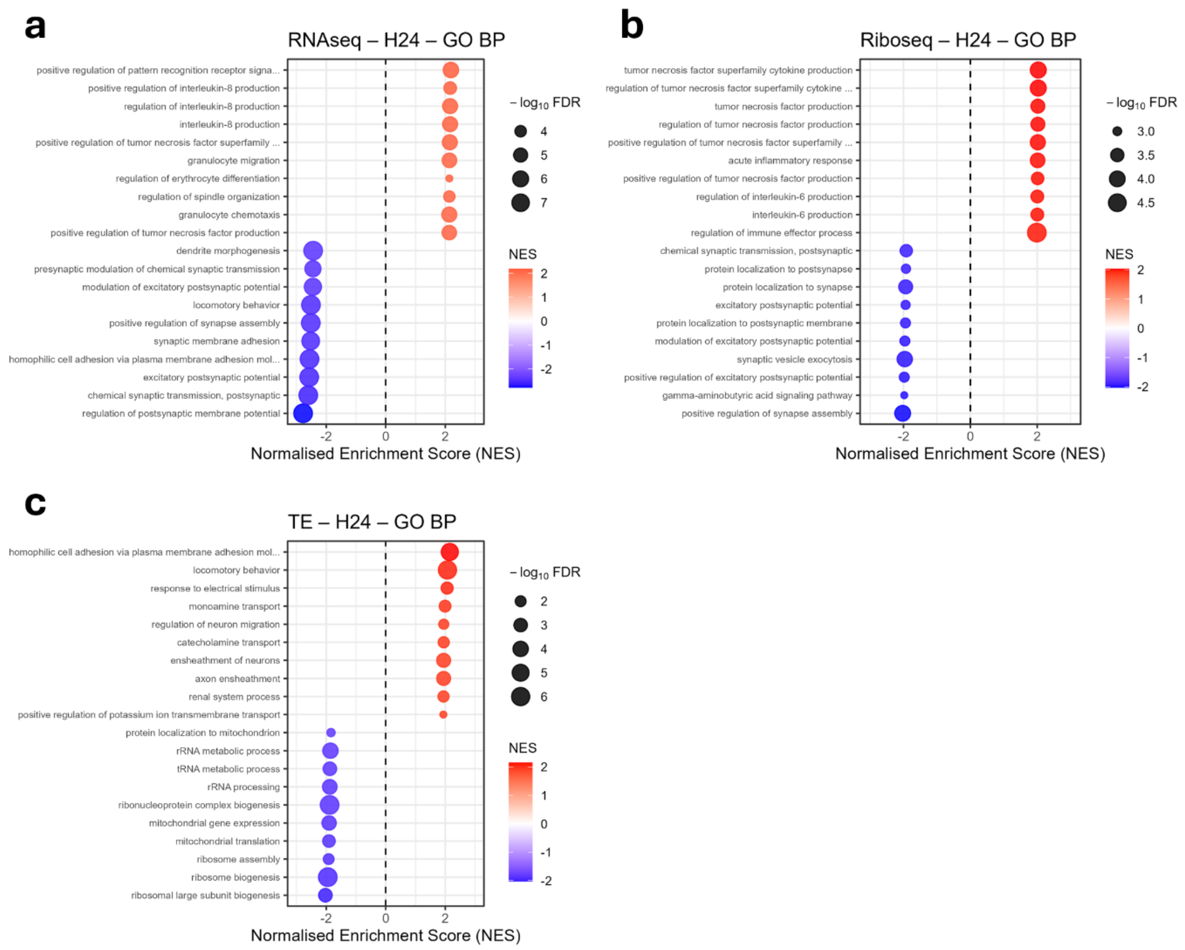


**Supplementary figure 2: a-c:** Pre-ranked GSEA analysis using GOBP gene set on different gene regulatory modes 24 hours after stroke.

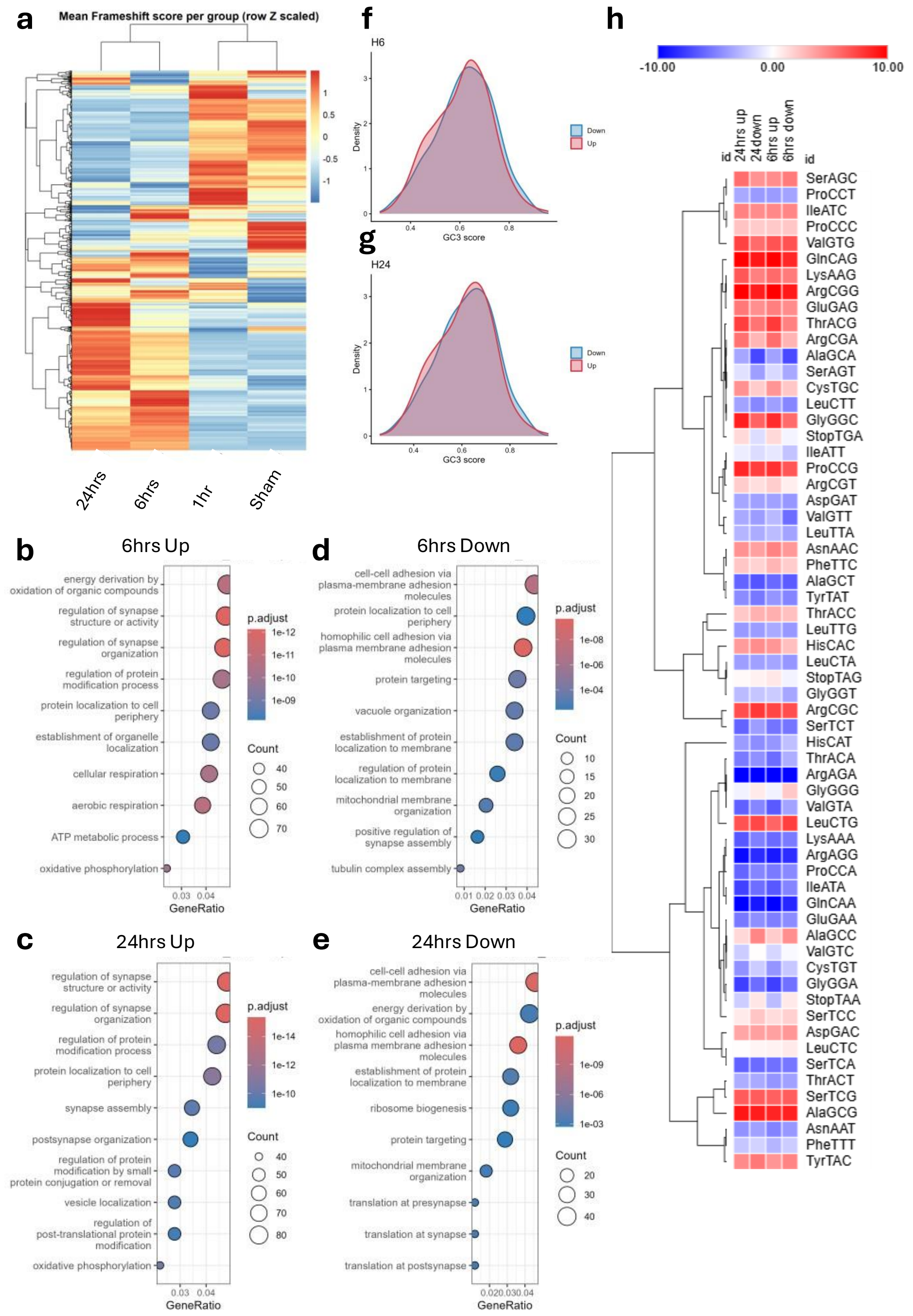


**Supplementary figure 3: a:** Heatmap of mean group frameshift (FS) scores. **b-e:** ORA GOBP analysis of pathway enrichment of genes showing differential frame usage (frameshifting changes) 6 and 24 hours after stroke. **f-g:** GC3 score density plot of up and downregulated genes in the frameshifting analysis 6 hours (**f**) and 24 hours (**g**) after stroke. **h:** isoacceptors codon frequencies analysis of up and downregulated genes in the frameshifting analysis 6 and 24 hours after stroke.

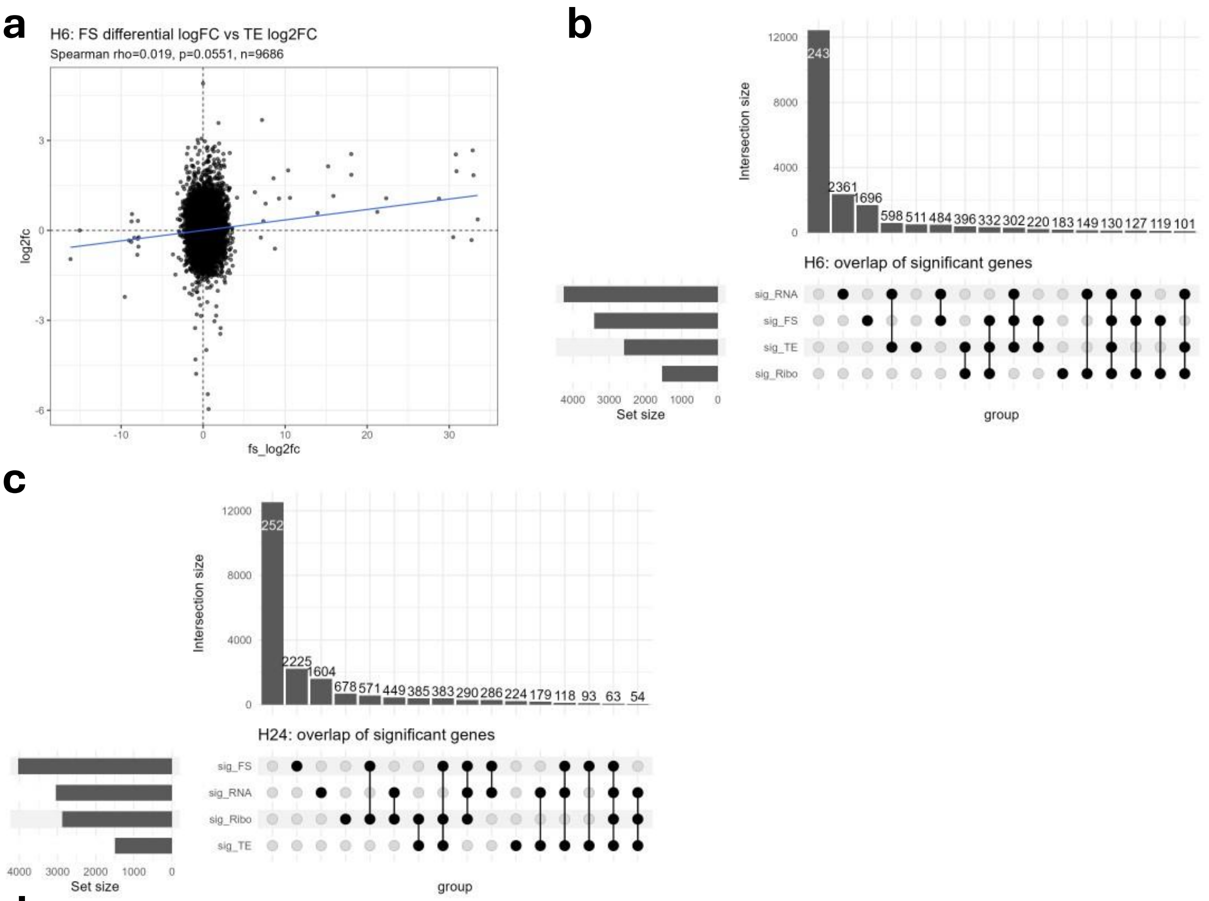


**d**

| condition | dataset | FS_class | Log2FC mean_class | log2FC mean_NoFS | Log2FC median_class | Log2FCmedian_NoFS | n_class | n_NoFS | ks_stat | p | padj |
|---|---|---|---|---|---|---|---|---|---|---|---|
| 6 hours | TE | FS+NMD | 0.123933263 | 0.012865372 | 0.117951379 | -0.008418106 | 168 | 11842 | 0.135337 | 0.004631 | 0.018523 |
| 6 hours | RNAseq | FS+NMD | -0.093993102 | 0.053033572 | -0.100153502 | 0.005354936 | 168 | 19485 | 0.200025 | 3.26E-06 | 1.30E-05 |
| 6 hours | Riboseq | FS+PTC_no_NMD | 0.157341896 | 0.03695304 | 0.14622808 | 0.001313246 | 28 | 11763 | 0.309053 | 0.009629 | 0.012839 |
| 6 hours | Riboseq | FS+SCRT | 0.094618031 | 0.03695304 | 0.031232898 | 0.001313246 | 92 | 11763 | 0.178821 | 0.005829 | 0.011658 |
| 6 hours | Riboseq | FS+NMD | 0.065453597 | 0.03695304 | 0.017910645 | 0.001313246 | 169 | 11763 | 0.133213 | 0.005407 | 0.011658 |
| 24 hours | RNAseq | FS+NMD | -0.064879515 | 0.10927161 | -0.072835511 | 0.005653478 | 215 | 19390 | 0.131543 | 0.001274 | 0.005095 |
| 24 hours | Riboseq | FS+SCRT | 0.116072305 | 0.070094616 | 0.113276632 | 0.005260412 | 105 | 11930 | 0.142689 | 0.028862 | 0.11545 |

**Supplementary figure 4: a:** Scatter plot showing Spearman's rank correlation between log2FC of genes in the FS analysis and in TE analysis at the 6 hours timepoint. **b-c:** Upset plots showing the overlap between significant genes in frameshifting analysis and gene regulatory modes (RNA-seq, Ribo-seq, and TE). The first column shows the total number of the genes used in the dataset. **d:** Table showing how frameshift impact influence gene expression and translation significantly at each time point. Komogrov-Smirnoff statistical test with Benjamin-Hochberg multiple test correction (FDR) was used for statistical analysis.

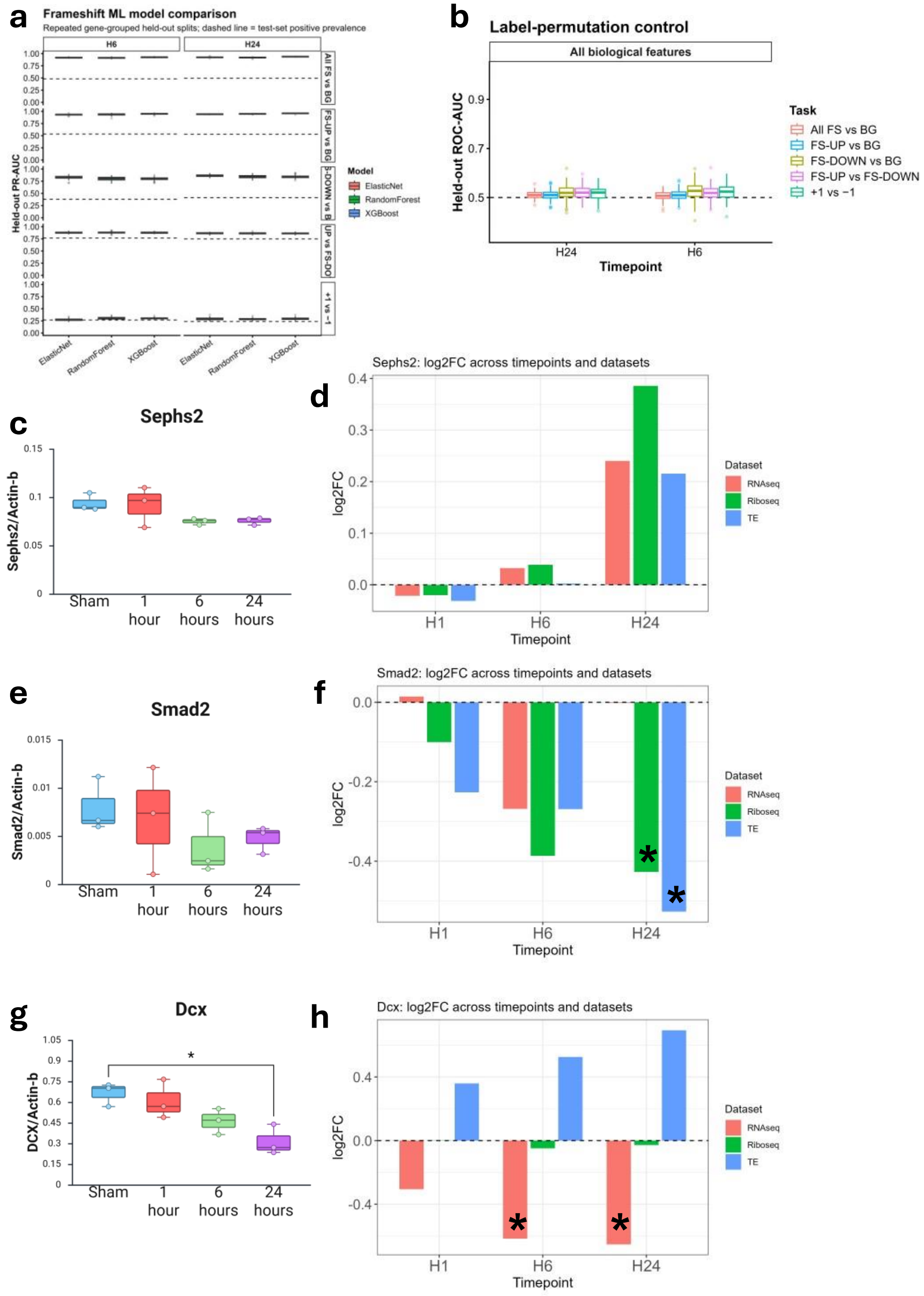


**Supplementary figure 5: a:** Held-out PR-AUC performance for elastic net, random forest, and XGBoost classifiers across repeated gene-grouped train/test splits, complementing the ROC-AUC analysis shown in Figure 4a. **b:** Label-permutation control for the repeated XGBoost models. Labels were permuted before model training while preserving the grouped-split structure. Model performance collapsed to near chance, indicating that the observed classification performance was not explained by generic overfitting or the split design. **c:** Supporting western blot quantification or replicate-level analysis for Sephs2 across sham and post-stroke timepoints. **d:** RNA-seq, Ribo-seq, and translational-efficiency log2 fold-change values for Sephs2 across post-stroke timepoints. Asterisks indicate FDR < 0.05. **e:** Supporting western blot quantification or replicate-level analysis for Smad2 across sham and post-stroke timepoints. **f:** RNA-seq, Ribo-seq, and translational-efficiency log2 fold-change values for Smad2 across post-stroke timepoints. Asterisks indicate FDR < 0.05. **g:** Supporting western blot quantification or replicate-level analysis for Dcx across sham and post-stroke timepoints. **h:** RNA-seq, Ribo-seq, and translational-efficiency log2 fold-change values for Dcx across post-stroke timepoints. Asterisks indicate FDR < 0.05.

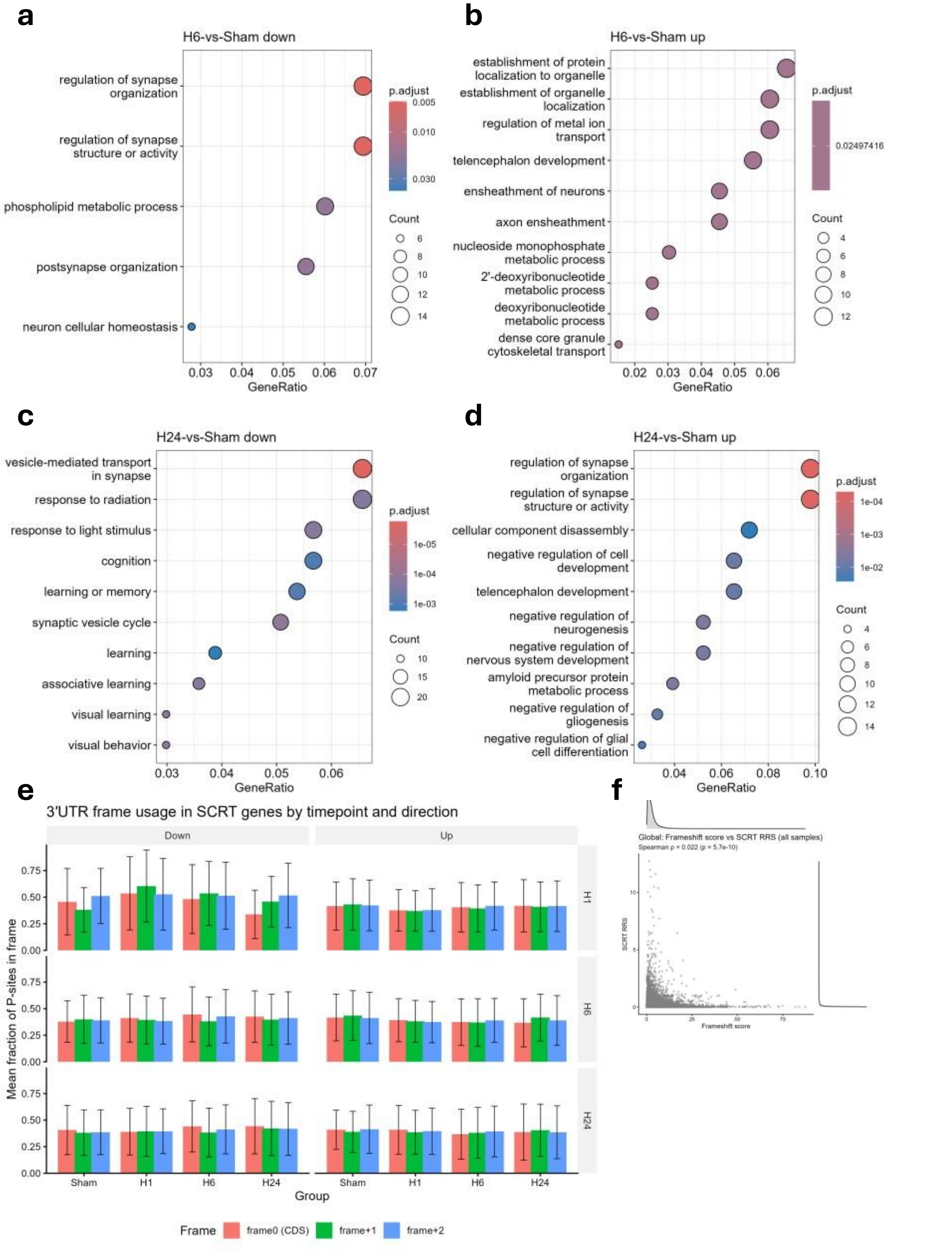


**Supplementary figure 6: a-d:** ORA GOBP analysis on up and downregulated genes in the SCRT analysis. **e:** RPFs frame in the 3'UTR. We selected significant genes at each time point and plotted the 3'UTR frames of their RPF reads across all time points. **f:** Spearman's rank correlation analysis of gene RRS scores and FS scores. **g:** eRF1 p-site density plot across the CDS 1 hour after stroke.

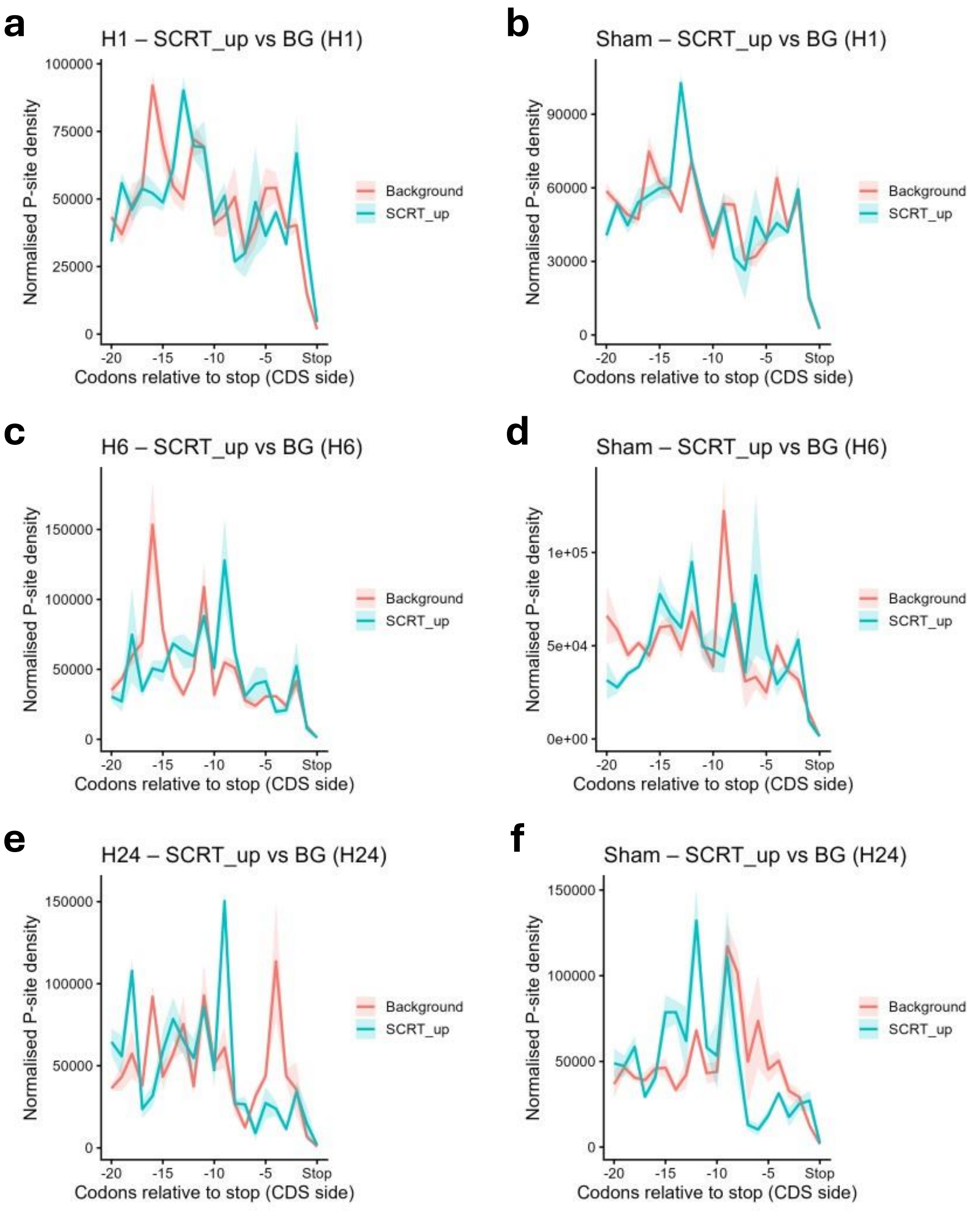


**Supplementary figure 7:** Pre-stop peaks in the upregulated SCRT genes at each time point compared to matched-coverage background genes in condition and sham. Significant upregulated genes were selected at 1- (**a-b**), 6- (**c-d**), and 24-hours (**e-f**) and their pre-stop P-site density was plotted compared to matched coverage background genes.

**a**

| group1 | group2 | p_value | median_MFE_g1 | median_MFE_g2 | Number_genes_g1 | Number_genes_g2 |
|---|---|---|---|---|---|---|
| Up | Background | 0.071935 | 20.9 | 22.4 | 197 | 4449 |
| Down | Background | 0.166202 | 23.2 | 22.4 | 223 | 4449 |
| Up | Down | 0.017816 | 20.9 | 23.2 | 197 | 223 |

**b**

| group1 | group2 | p_value | median_MFE_g1 | median_MFE_g2 | Number_genes_g1 | Number_genes_g2 |
|---|---|---|---|---|---|---|
| Up | Background | 0.301669 | 24.4 | 22.3 | 153 | 4369 |
| Down | Background | 0.168236 | 23.1 | 22.3 | 347 | 4369 |
| Up | Down | 0.906162 | 24.4 | 23.1 | 153 | 347 |

**c**

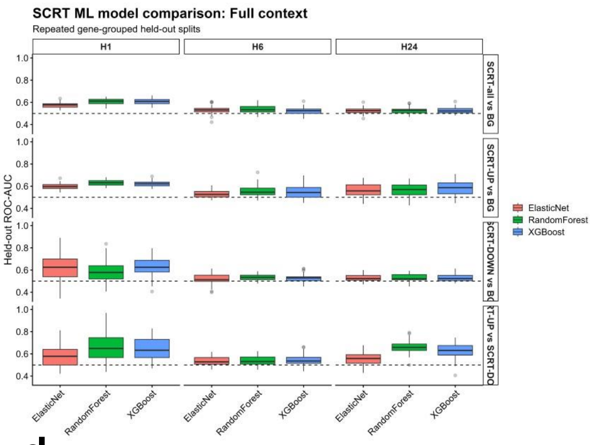


**d**

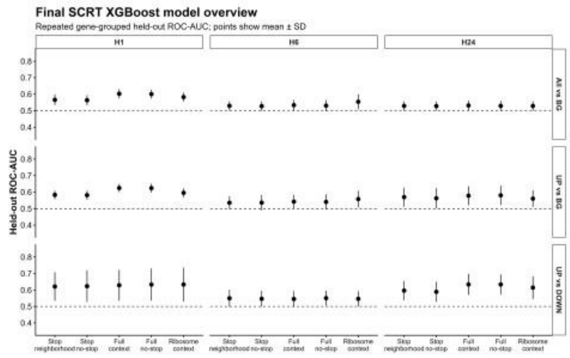


**e**

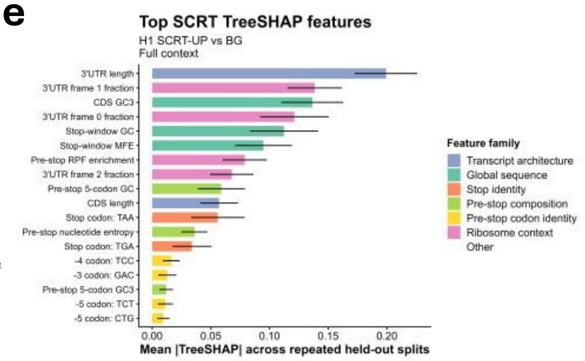


**f**

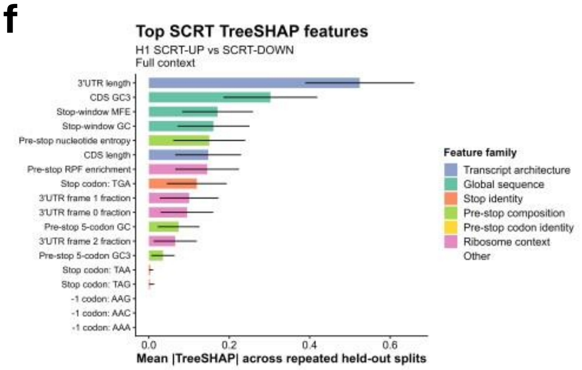


**g**

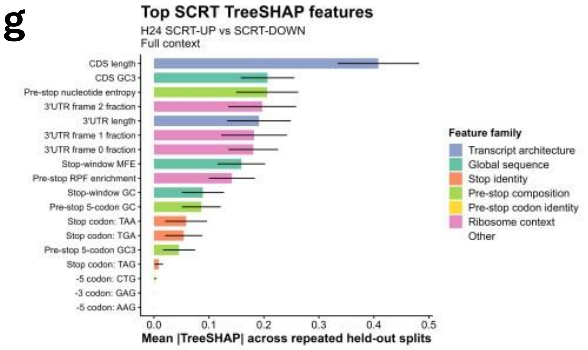


**h**

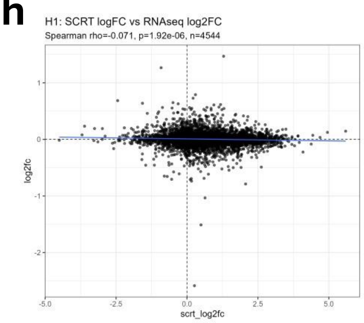


**i**

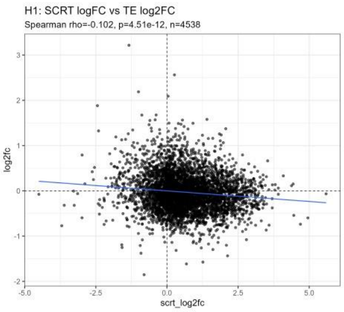


**j**

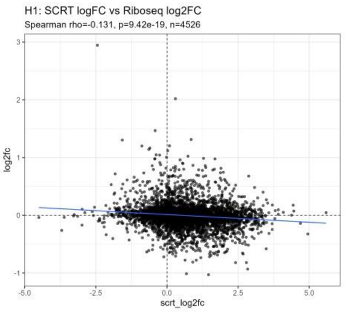

**Supplementary figure 8: a-b:** Tables showing Wilcoxon test results comparing MFE of the first 100 3'UTR nucleotides downstream of the stop codon of upregulated or downregulated genes in SCRT analysis 6- (**a**) or 24-hours (**b**) after stroke. **c:** Model-comparison analysis for SCRT classification using elastic-net, random forest, and XGBoost classifiers across timepoints and SCRT tasks. Models were evaluated using repeated gene-grouped held-out splits. **d:** Final XGBoost feature-panel sensitivity analysis across timepoints and SCRT classification tasks. Points show mean held-out ROC-AUC ± SD across repeated gene-grouped splits for stop-neighborhood, stop-neighborhood without stop identity, full-context, full-context without stop identity, and ribosome-context-only feature panels. **e:** Top TreeSHAP features from the H1 SCRT-UP versus background full-context model including stop codon identity. **f:** Top TreeSHAP features from the H1 SCRT-UP versus SCRT-DOWN full-context model. **g:** Top TreeSHAP features from the H24 SCRT-UP versus SCRT-DOWN full-context model. **h-j:** Spearman's rank correlation analysis between gene SCRT log2FC and log2FC values from RNA-seq (**h**), TE (**i**), Ribo-seq (**j**) datasets.

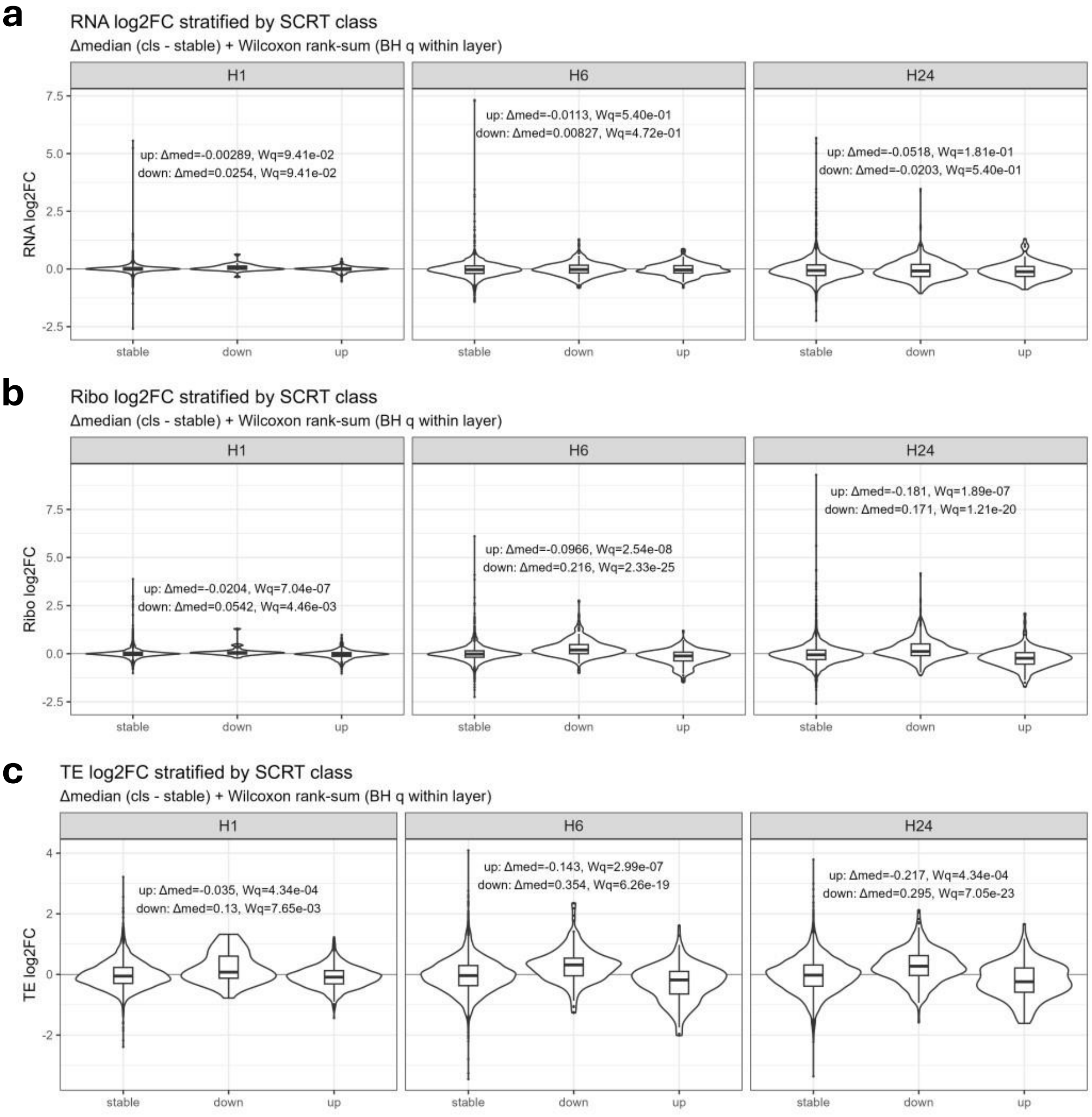


**Supplementary figure 9:** Violin plot showing log2FC of genes with SCRT changes across modes and time points. Wilcox rank sum test with BH correction results are shown with delta median log2FC values compared to stable (change in median log2FC compared to stable genes (i.e. no significant SCRT changes compared to Sham).

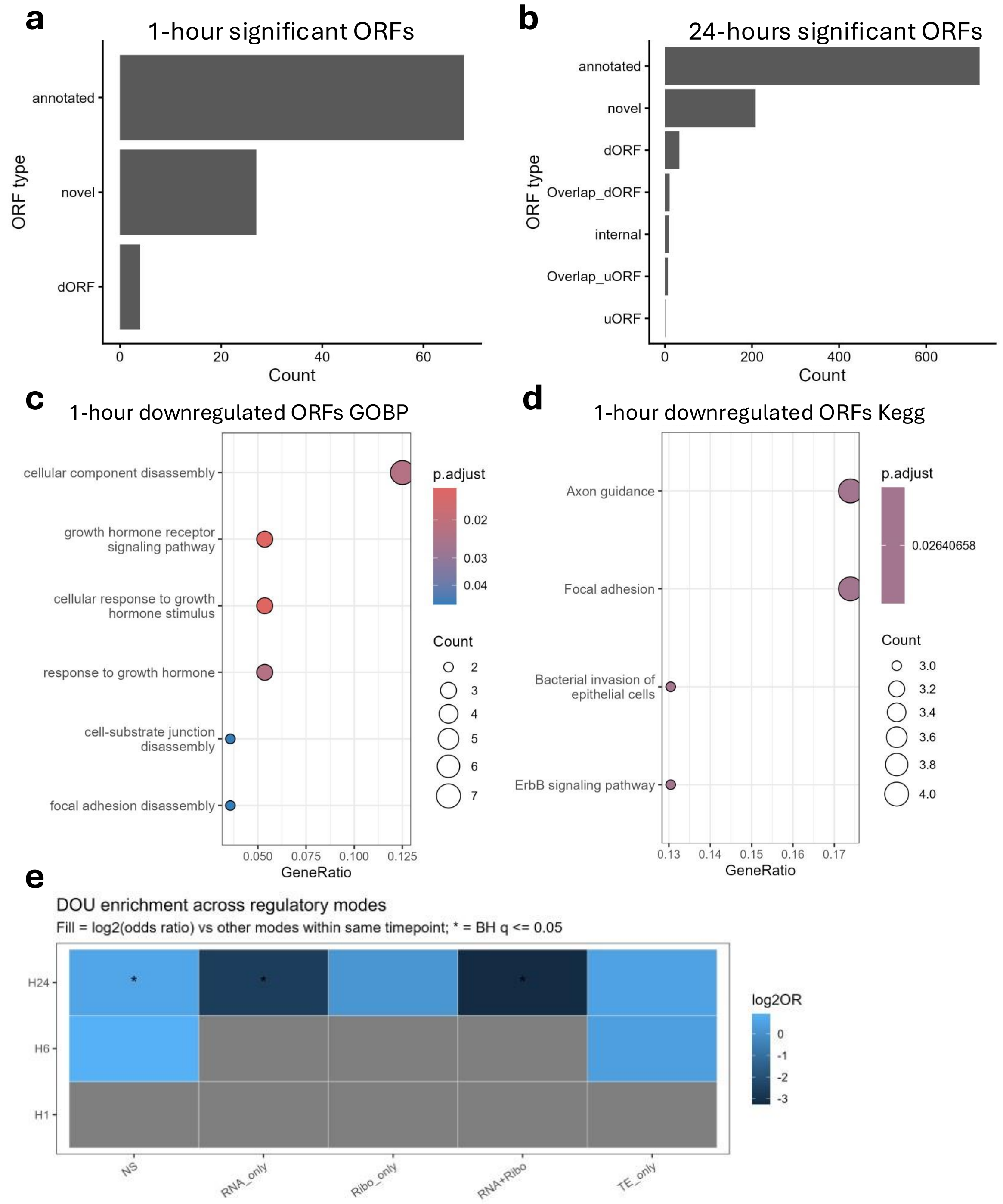


**Supplementary figure 10: a-b:** Bar plots showing the ORF annotation of significant differentially expressed ORFs 1- and 24-hours after stroke (related to figure 7c and 7e). **c-d**: ORA pathway analysis of downregulated ORF genes 1 hour after stroke. **e:** Analysis of potential gene regulatory mode linked to DOU using Fisher's exact test with BH multiple test correction.

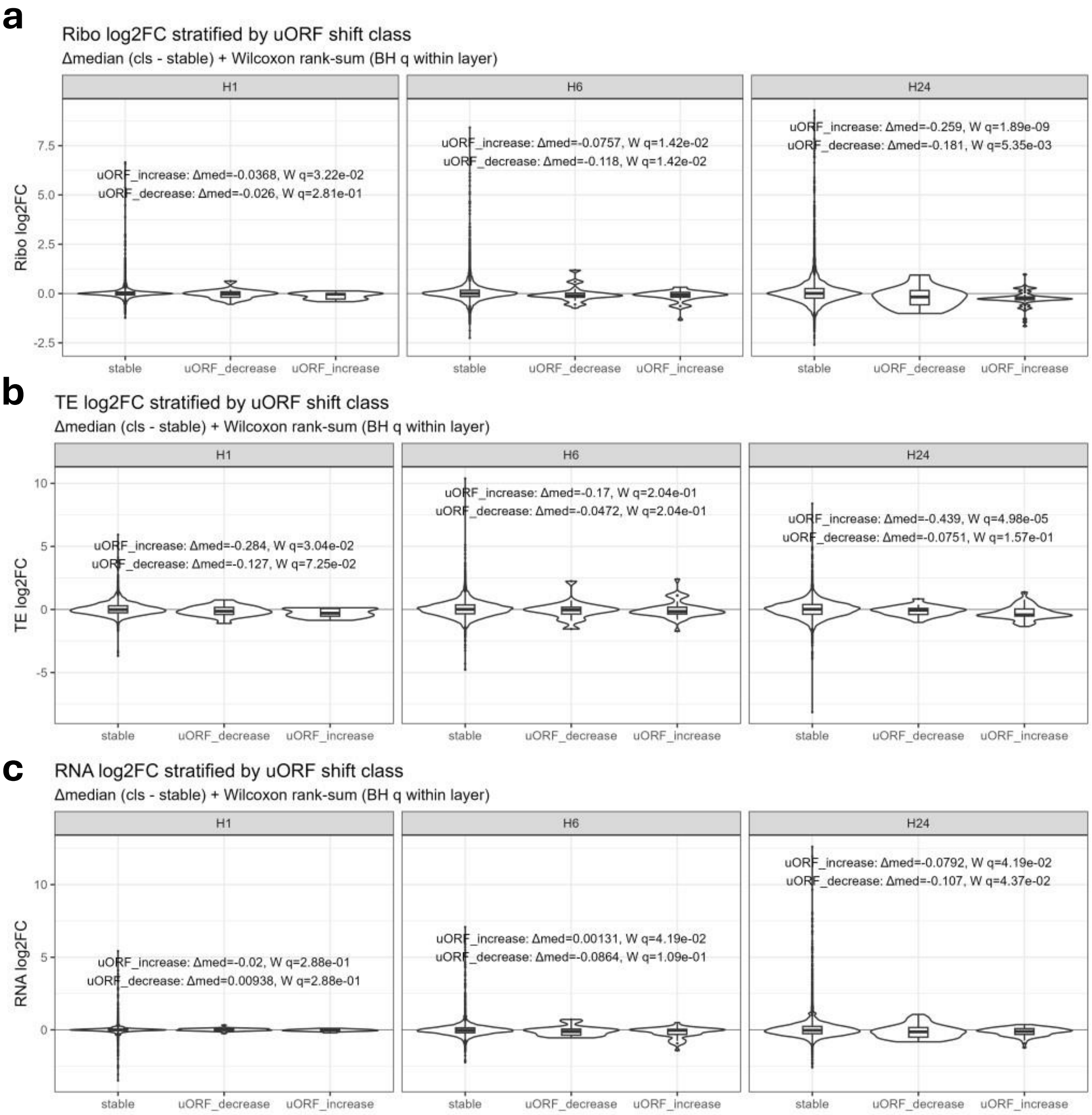


**Supplementary figure 11:** Violin plot showing log2FC of genes with uORF usage changes across modes and time points. Wilcox rank sum test with BH correction results are shown with delta median log2FC values compared to stable genes (no uORF change)

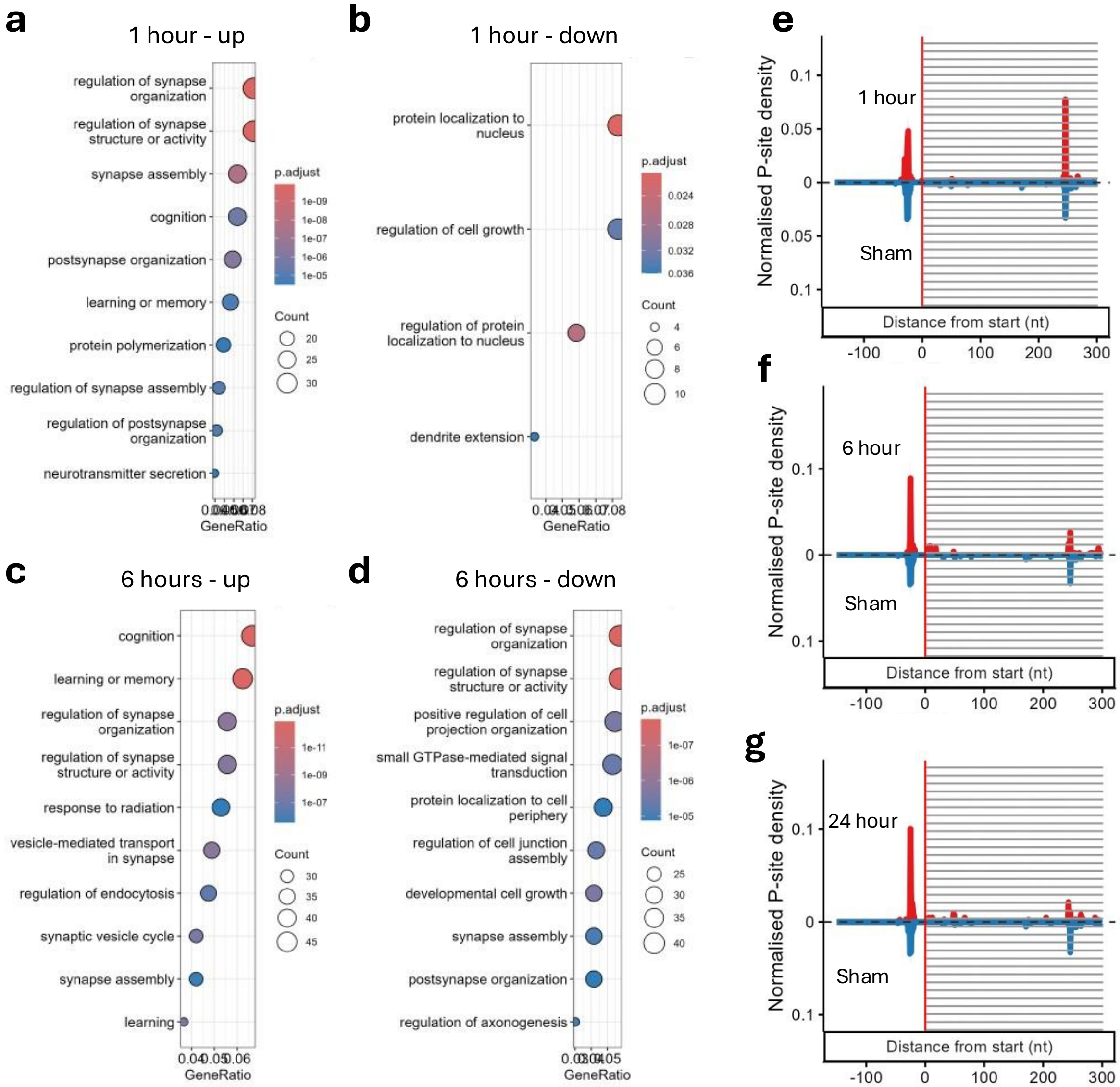


**Supplementary figure 12: a-d:** ORA GOBP analysis of genes significant in upstream translation analysis at 1- and 6-hour timepoints. **e-g:** Normalized P-site density metagene plots showing the pre-start increase in 5'UTR P-site peaks of Smad2 gene over time.

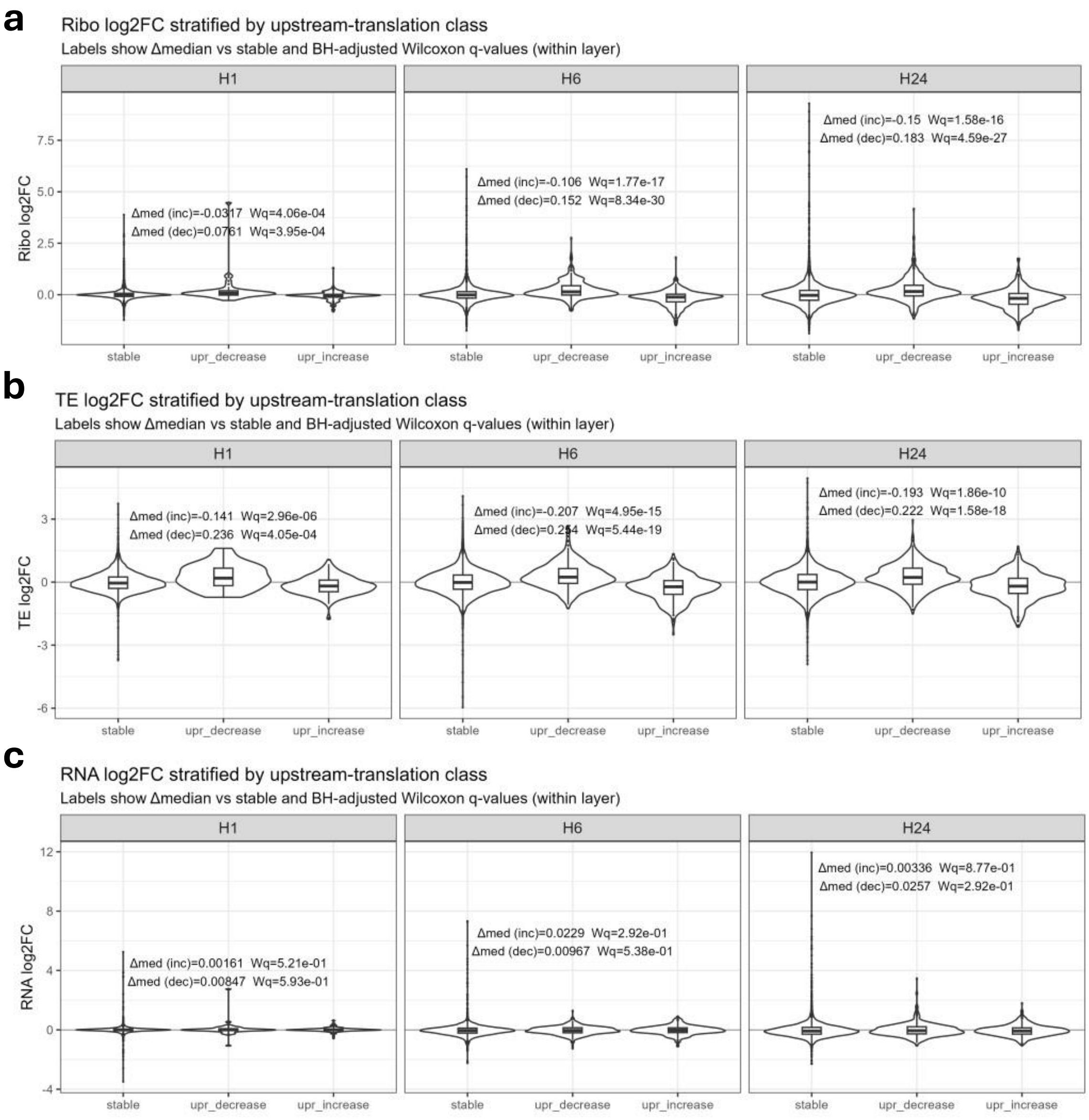


**Supplementary figure 13:** Violin plot showing log2FC of genes with Upstream translation usage changes across modes and time points. Wilcox rank sum test with BH correction results are shown with delta median log2FC values compared to stable genes (no change in 5'UTR usage compared to Sham controls).
